# Untangling the diversity of Trypanosomes infecting Colombian amphibians: morphometric and molecular insights

**DOI:** 10.1101/2025.10.25.684581

**Authors:** Angélica T. Ospina-Rios, Angie D. Gonzalez Galindo, Andrés F. Aponte-Gutiérrez, Carolina M. Vargas-León, Nubia E. Matta

**Author notes:** Corresponding author: Nubia E. Matta.

## Abstract

Amphibians, though widely distributed and diverse, are increasingly threatened by environmental changes and emerging diseases. Among the lesser-known yet prevalent pathogens affecting them are *Trypanosoma* parasites, which remain poorly studied, particularly in terms of taxonomy and molecular diversity. In Colombia, one of the most amphibian-rich countries globally, no prior studies had addressed the molecular identification of *Trypanosoma* species in anurans. This study aimed to strengthen the taxonomy of these parasites by integrating morphological, morphometric, and molecular analyses. A total of 434 amphibian samples from the GERPH biological collection were examined microscopically, with 92 testing positive for *Trypanosoma*. Thirteen morphometric and eleven categorical variables were used to characterize the parasites, leading to the identification of twelve distinct morphotypes through qualitative classification and multivariate analysis. Of the positive samples, 48.9% exhibited more than one morphotype, with morphotypes I, II, and III being the most widespread and generalist. To explore molecular diversity, DNA from 76 individuals positive by microscopy was analyzed using PCR targeting the 18S rRNA gene, yielding nine molecular lineages in 38 single-infection cases. Five morphotypes were successfully linked to molecular lineages, enabling barcode associations that facilitated the description of a new species, the expansion of the known range of *Trypanosoma tungarae*, and the identification of a potential species complex. This study highlights the importance of integrating morphological and molecular tools to elucidate parasite taxonomy and host-parasite relationships, underscoring the need for further research into vector ecology, molecular barcoding, and *in vivo* isolation methods to understand *Trypanosoma* diversity in amphibians.

## 1. Introduction

The genus *Trypanosoma* (Gruby, 1843) comprises a widely distributed group of protozoan parasites that infect various vertebrates, including amphibians. These parasites are transmitted through hematophagous vectors such as dipterans, leeches, and sandflies, or ingestion of infected vectors or contaminated feces (Ferreira et al., 2008; Hayes et al., 2014). In humans, these parasites cause Chagas disease (*Trypanosoma cruzi*) or African sleeping sickness (*Trypanosoma brucei*) (Truc et al., 2013). These diseases are characterized by severe pathological signs such as anemia, fever, lethargy, emaciation, and, in extreme cases, death. Despite the extensive research on *Trypanosoma* in humans and livestock, the impact of these parasites on wildlife, particularly amphibians, remains neglected (Spodareva et al., 2018).

Amphibians, such as frogs and toads (Anura), salamanders (Caudata), and caecilians (Gymnophiona), are especially vulnerable to emerging pathogens. As one of the most biodiverse countries in the world, Colombia is home to around 902 amphibian species, 359 of which are listed as threatened (Acosta-Galvis, 2023). Unfortunately, amphibians are also the most endangered group of vertebrates, facing numerous threats, including climate change, pollution, habitat loss, and emerging diseases like chytridiomycosis, caused by the fungus *Batrachochytrium dendrobatidis* (*Bd*), which has contributed to the extinction of approximately 90 species (Herczeg et al., 2021; Sewell et al., 2021). Their anatomy, including thin skin that facilitates gas exchange, makes amphibians particularly susceptible to infections (Grogan et al., 2018). While the effects of well-known pathogens, such as *Bd,* are well-documented, information on other parasites, such as *Trypanosoma* spp., remains limited. However, some studies characterizing hemoparasites in amphibians have identified *Trypanosoma* as one of the most prevalent (Barta & Desser, 1984; Netherlands et al., 2015; González et al., 2021; Rodrigues et al., 2019), and it is still unknown whether they contribute to population declines (Bernal & Pinto, 2016).

Gabriel Valentin made the first description of a trypanosomatid in 1841, grouping several flagellated forms under the name Trypanoplasma. Two years later, David Gruby formally proposed the genus *Trypanosoma* and described *Trypanosoma sanguinis,* identifying for the first time a species of this genus in an amphibian blood sample (Gruby, 1843). The characterization of *Trypanosoma* in anurans was initially based only on morphological and morphometric characteristics, which allows describing species such as *T. rotatorium* (Mayer, 1843), *T. chattoni* (Mathis & Léger, 1911), *T. mega*, *T. neveulemairei* (Brumpt, 1928), *T. ranarum*, and *T. fallisi* (Martin & Desser, 1990).

However, *Trypanosoma* taxonomy is complicated by the parasites’ pleomorphic nature, where morphology varies across life stages, species, and hosts, making identification challenging (Hamilton et al., 2004; Spodareva et al., 2018). Due to their pleomorphic nature, these parasites can exhibit multiple morphologies within the same species, making exclusive diagnosis by morphology difficult and underscoring the need for molecular tools for accurate characterization (Bardsley & Harmsen, 1973; Martin et al., 2002). The molecular markers commonly used are the 18S rRNA gene (Noyes et al., 1999) and, in some cases, the GAPDH marker (Hamilton et al., 2004).

The development of molecular tools has allowed for more accurate identification and characterization of *Trypanosoma* species, shedding light on their phylogenetic relationships and clarifying species complexes (Martin et al., 2002; Bartlett-Healy et al., 2009; Ferreira et al., 2007; Lemos et al., 2008). Molecular analyses using genetic markers such as 18S rRNA and gGAPDH genes have revealed that *Trypanosoma* species can be grouped into clades based on their vertebrate hosts. Notably, an aquatic clade has been identified, which includes trypanosomes infecting amphibians, reptiles, and fish, suggesting a common evolutionary origin for these species (Spodareva et al., 2018). Furthermore, molecular studies on *Trypanosoma* in their vectors have shown that these parasites are distributed across various clades of vertebrates, further complicating the classification of different lineages (Hamilton et al., 2004; da S. Ferreira, 2015; Spodareva et al., 2018).

Taking into account the high richness of amphibians in Colombia and the high prevalence of *Trypanosoma* infections in this taxonomic group. We will likely discover new host-parasite associations in this research. The vast amphibian diversity and the number of *Trypanosoma* species potentially infecting them present an opportunity for further investigation into the molecular diversity of these parasites. Research on *Trypanosoma* infections in amphibians could provide insights into their ecological impact, such as how these infections might influence behavior, reproduction, and competition within anuran populations. Additionally, the potential for amphibians to serve as reservoirs for emerging infectious diseases highlights the need for ongoing monitoring of *Trypanosoma* in wildlife (Truc et al., 2013). The study of *Trypanosoma* in amphibians can also contribute to broader efforts to understand how parasites from wildlife species may spill over to human populations, as has been documented in other cases of zoonotic diseases (Jansen et al., 2018).

Using samples from the Grupo Estudio Relación Parásito Hospedero (GERPH) Biological Collection at the Universidad Nacional de Colombia, this study aims to characterize *Trypanosoma* parasites infecting anurans through an integrated approach combining morphometric and molecular analyses.

## 2. Materials and methods

### 2.1 Samples

The GERPH Biological Collection of the Department of Biology of the Universidad Nacional de Colombia has in its repository blood samples from 434 frogs and toads collected between 2018 and 2022 in ten localities of Colombia: in the Trans-Andean area: 1. Magdalena (Santa Marta), 2. Magdalena (Pueblo Viejo), 3. Santander (San Gil), 4. Cundinamarca (Tibacuy), 5. Nariño (Tumaco). In the Cis-Andean area: 6. Casanare (Trinidad), 7. Casanare (El Yopal), 8. Cundinamarca (Medina), 9. Guaviare (San José del Guaviare), and 10. Caquetá (Solano-Araracuara) (Suppl. Fig. 13, Suppl. Table S1). The samples used in this study correspond to blood smears obtained by puncturing the maxillary or abdominal veins (Forzán and Wood, 2013). The smears were fixed in methanol for 5 minutes and stained with 4% Giemsa.

### 2.2 Microscopic diagnosis

A total of 434 samples were examined (Suppl. Table S1). For trypanosome diagnosis, a complete screening of blood smears per sample was performed by visualizing the entire slide at magnification of 100X or 400X, and photographs were taken at 1000X using an Olympus BX43 light microscope (Olympus Corporation, Tokyo, Japan) with a built-in CellSens camera (Olympus Corporation). Parasitemia was calculated as the average trypanosomatids found in 2-3 blood smears.

### 2.3 Morphological and morphometric characterization

Measurements relative to five key structures (e.i. overall structure, nucleus, kinetoplast, undulating membrane, and flagellum) were taken by using ImageJ software Version 1.54i RRID: SCR_003070 (National Institute of Health, Bethesda, Maryland, United States) (Schneider et al. 2012) as follows (Fig. 1): ***Overall structure:*** Shape, arrangement, total length without flagellum (L), width at the level of the nucleus - without membrane (W), anterior border shape, posterior border shape, cytoplasm texture, presence/absence of vacuoles, total area (A). ***Nucleus:*** shape, distance anterior edge-nucleus (AN), distance posterior edge-nucleus (PN), major diameter of the nucleus (DN), minor diameter of the nucleus (dN), nuclear area (nA). ***Kinetoplast:*** shape, kinetoplast length (KL), distance from kinetoplast to nucleus (KN), distance from kinetoplast to posterior edge (PK). ***Undulating membrane:*** Presence/absence of membrane, visible axoneme, wavenumber, undulating membrane width (MW). ***Flagellum:*** length of free flagellum (FF). Thirty-three categories assigned to eleven qualitative features are shown in the Supplement. Table S2.

**Fig. 1.**
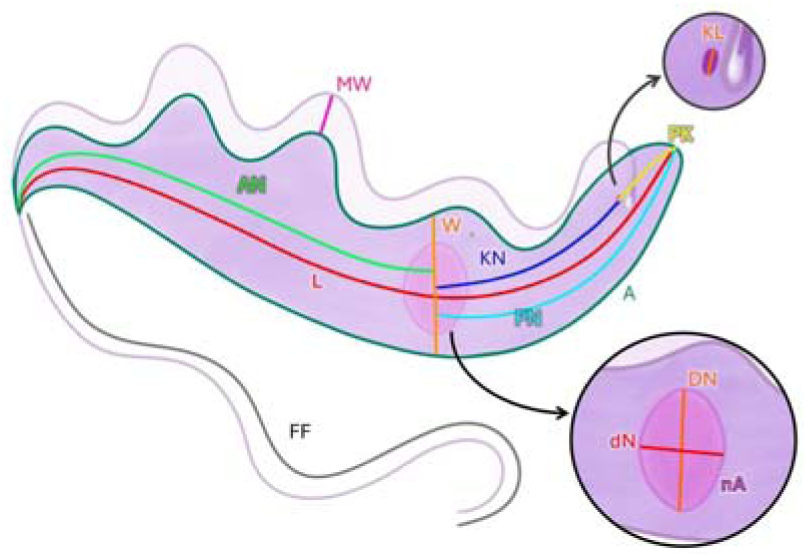
Morphometric measurements taken for *Trypanosoma* spp. Morphological characterization. Total length without flagellum (L), parasite width at the level of the nucleus (W), and total area (A). To nucleus: distance anterior edge-nucleus (AN), distance posterior edge-nucleus (PN), major diameter of the nucleus (DN), Minor diameter of the nucleus (dN), nuclear area (nA). To kinetoplast: kinetoplast length (KL), distance from kinetoplast to nucleus (KN), distance from kinetoplast to posterior edge (PK). Undulating membrane width (MW). Flagellum: length of free flagellum (FF).

### 2.4 Statistical analysis

Only morphotypes with at least 30 measured photographs were analyzed. The normal distribution of the 13 quantitative variables was evaluated through a Shapiro-Wilk normality test using Past Version 4.02 RRID: SCR_019129 (Hammer et al. 2001). A Principal Component Analysis (PCA) was performed to graphically observe the components and their variables that contribute most to the variation of morphotypes. Two multivariable non-parametric tests were used to determine whether morphotypes can be distinguished by morphometry: Analysis of similarity ANOSIM (Clarke and Green, 1988; Clarke, 1993) and Permutational multivariate analysis of variance PERMANOVA. For both analyses, 1000 permutations were used, and for pair-comparison, the significance was calculated using Bonferroni-corrected p-values. The variables analyzed were W, DN, PK, KN, KL, AN, PN, and MW). Finally, a cluster analysis using Gower distance was done with 13 morphometric and 11 categorical variables, and the results were presented as a cladogram. An individual of morphotype XII was used as the root, as it is morphologically distinct.

### 2.5 DNA extraction, PCR amplification, and sequencing

DNA extractions were performed using the Quick-DNATM Miniprep Plus extraction kit (Zymo Research) or the Sigma kit (Sigma-Aldrich, Missouri, United States). DNA quality was evaluated using a NanoDrop One (Thermo Fisher Scientific, Waltham, Massachusetts, United States) to determine the 260/230 and 260/280 ratios. Nested PCR was performed on 76 samples (based on the available blood or tissue samples) using a protocol adapted from Jordaan et al. (2023a, 2023b), with minor modifications, to amplify the *Trypanosoma* 18S gene. The reaction mix used for the two successive rounds of amplification contained: 1X PCR buffer, 3 mM MgClL, 1.28 mM of each dNTP, 0.6 µM of each primer, 3.1 U Taq DNA polymerase, and 50 ng of template DNA, on a final volume of 25 μl. For the first round, primers SLF (5′-GCTTGTTTCAAGGACTTAGC-3′) and S762.2 (5′-GACTTTTGCTTCCTCTAATG-3′) were used under the following cycling conditions: one cycle with an initial denaturation at 95 °C for 5 min, annealing at 50°C for 2 min, and extension at 72°C for 4 min, followed by 35 cycles of denaturation at 94 °C for 30 s, annealing at 54 °C for 30 s, and extension at 72 °C for 2 min and 20s, with a final extension at 72 °C for 7 min. For the second round, primers S825 (5′-ACCGTTTCGGCTTTTGTTGG-3′) and SLIR (5′-ACATTGTAGTGCGCGTGTC-3′) were used under the following cycling conditions: initial denaturation at 95°C for 3 min, followed by 35 cycles of denaturation at 95°C for 30 s, annealing at 60°C for 30 s, and an extension step at 72°C for 1 min with a final extension at 72°C for 7 minutes.

Amplified products were visualized on a 1.5% agarose gel stained with SYBR Safe, and products purified from the gel using the Wizard® SV Gel and PCR Clean-Up System band purification kit. DNA sequences were obtained by Sanger sequencing at Macrogen Inc. (South Korea).

### 2.6 Phylogenetic and genetic distance analyses

FASTA sequences were processed for quality control using Sequencer v4.1.4 (Gene Codes Corporation, Michigan, United States). Low-quality bases and primer sequences were manually removed using BioEdit (Hall, 1999). Sequences retaining at least 80% of the original read length were used for downstream analysis. A total of 65 sequences were used for the phylogenetic analysis, including 57 sequences retrieved from the NCBI GenBank database, and eight lineages obtained in this study (Suppl. Table S3, Suppl Fig. 14). All the sequences were aligned using MAFFT V7 (Katoh & Standley, 2013) default settings, and maximum likelihood analysis (ML) analysis was performed using IQ-TREE software (Nguyen et al., 2015; Trifinopoulos et al., 2016; Hoang et al., 2018) following the (TIM3e+I+G4) evolutionary model suggested by Jmodeltest (Spodareva et al., 2018). Genetic distances were calculated using MEGA V11 (Molecular Evolutionary Genetics Analysis) (Tamura et al., 2021). The number of base substitutions per site was estimated, and pairwise distances were calculated using the Kimura 2-parameter substitution model

## 3. Results

### 3.1 Microscopic diagnosis

Four hundred thirty-four samples were analyzed, from 56 amphibian species across eight taxonomic families. Infection with *Trypanosoma* spp. was detected in 21.2% (92 out of 434) of the samples by microscopic examination. Of the 92 positive samples, 76 had available material (blood smears) for posterior analysis (Suppl. Table S3).

The positive samples were taken from 25 species belonging to six (6) taxonomic families sampled in six localities (Suppl. Table S3). The most abundant species sampled were *Rhinella horribilis* (122), *Rhinella marin*a (55), *Boana platanera* (29), and *Leptodactylus fuscus* (29). The most common infected species were *Leptodactylus latrans* (19), *Boana platanera* (15), and *Leptodactylus insularum* (7). *L. latrans*, *B. platanera*, and *Boana lanciformis* showed the lowest parasitemia (1), while *Scinax ruber* (184) and *Boana boans* (84) had the highest parasitemia (Suppl. Table S3).

### 3.2 Morphological and morphometric characterization

The detailed descriptions of each morphotype identified are shown below (Table 1).

**Table 1.**
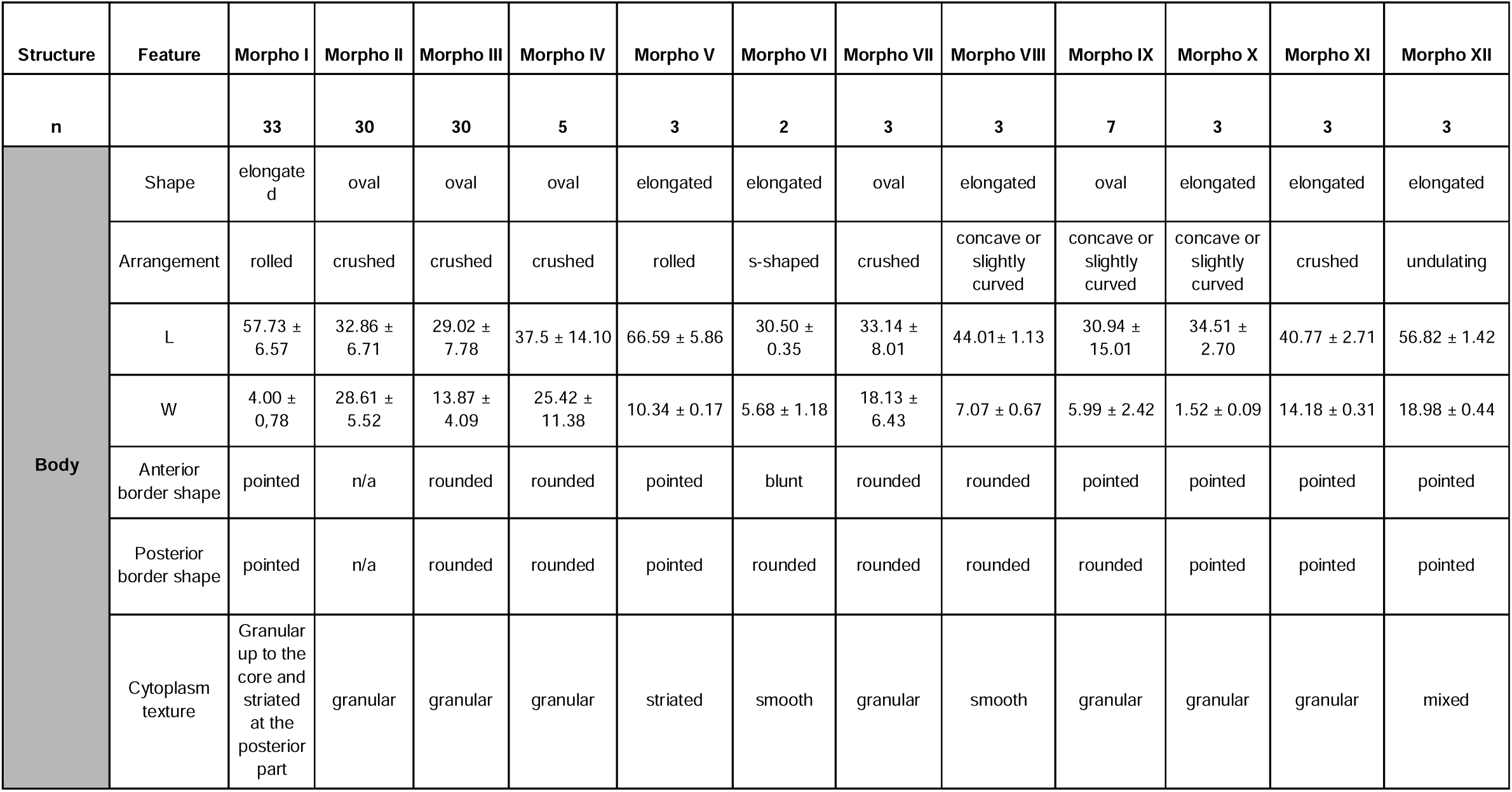
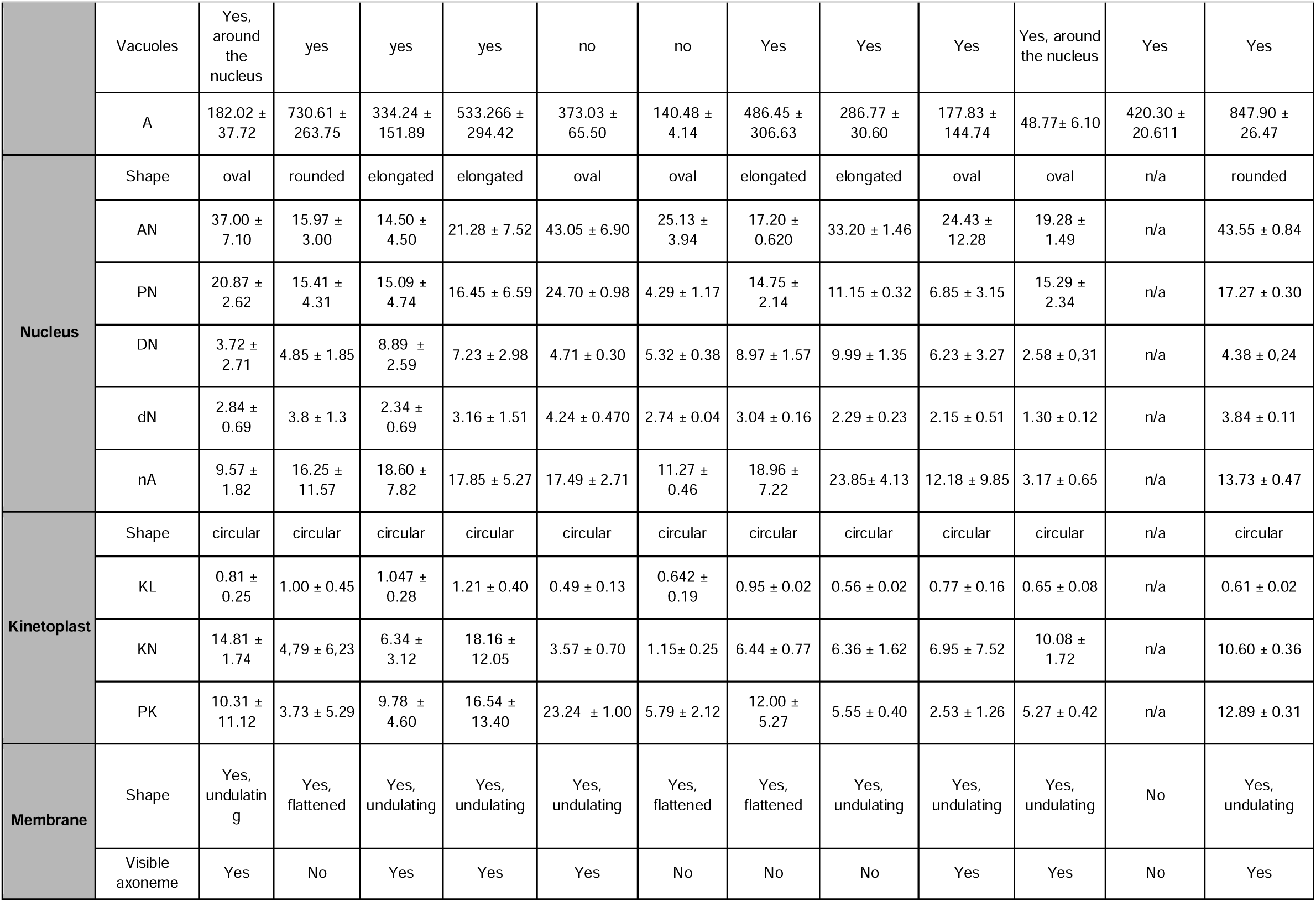
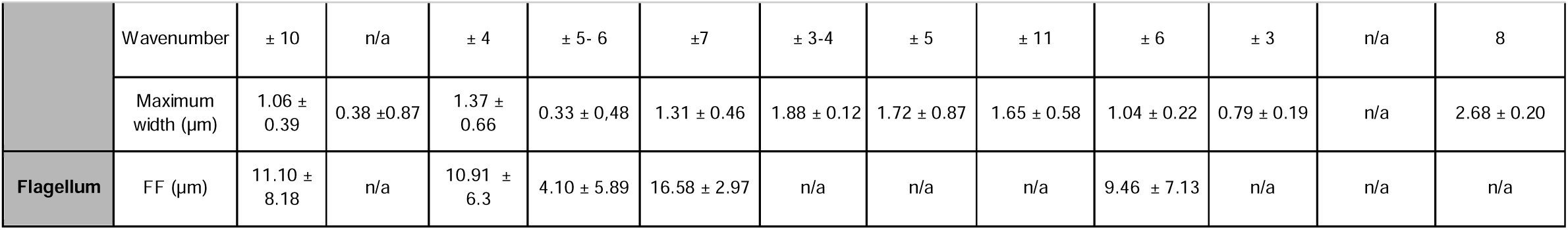
Average morphological and morphometric parameters of the twelve morphotypes of *Trypanosoma* spp. found in the amphibian sample analyzed. To overall structure: Shape, arrangement, total length without flagellum (L), width at the level of the nucleus - without membrane (W), anterior border shape, posterior border shape, cytoplasm texture, presence/absence of vacuoles, total area (A). To nucleus: shape, distance anterior edge-nucleus (AN), distance posterior edge-nucleus (PN), major diameter of the nucleus (DN), Minor diameter of the nucleus (dN), nuclear area (nA). To kinetoplast: shape, kinetoplast length (KL), distance from kinetoplast to nucleus (KN), distance from kinetoplast to posterior edge (PK). Undulating membrane: Presence/absence of membrane, visible axoneme, wavenumber, undulating membrane width (MW). Flagellum: length of free flagellum (FF). All measurements are represented in micrometers (µm) and areas in µm^2^.

#### Morphotype I

Found in *Leptodactylus latrans*, *Leptodactylus validus,* and *Pristimantis* sp. *Trypanosoma* elongated (57.73 µm ± 6.57), coiled on itself or S-shaped, with anterior and posterior edges pointed. It has a conspicuous nucleus (3.72 µm ± 2.71 x 2.84 µm ± 0.69) with an oval-rectangular shape that occupies the entire width of the parasite and is located in the middle posterior part. The kinetoplast is visible (0.81 µm ± 0.25) and is located 10.31 µm ± 11.12 from the posterior edge. Several vacuoles surround the nucleus, extending from the middle to the posterior region of the cytoplasm, while the anterior part of the cytoplasm is striated. The undulating membrane begins near the kinetoplast and extends throughout the parasite’s body, forming distinct undulations. The flagellum may or may not be visible; when visible, its length is approximately 1/5 of the parasite (11.10 µm ± 8.18) in length (Fig. 2-I, Suppl. Fig. 1).

#### Morphotype II

Found in *Rhinella horribilis*, *Leptodactylus latrans, Leptodactylus validus, Scinax rostratus, Leptodactylus insularum, Boana platanera, Boana boans, Scinax aff. ruber, Trachycephalus typhonius, Leptodactylus colombiensis, Lithobates palmipes, Smilisca phaeota, Leptodactylus fuscus, Hypsiboas maculateralis. Trypanosoma* with a flattened form (average length and width of 32.86 µm ± 6.71 x 28.61 µm ± 5.52) that tends to be rounded with defined edges. The nucleus is round (4.85 µm ± 1.85 x 3.8µm ± 1.3) and is located in the center of the parasite (PN: 15.41 µm ± 4.31, AN: 15.97 µm ± 3.00), and the nucleolus is not visible. The kinetoplast (1.00 µm ± 0.45) is located adjacent to or above the nucleus, and a short region of the undulating membrane can be observed coming out of the kinetoplast. The membrane rarely exceeds the length of the nucleus. The cytoplasm can be either granular or with well-defined striations (42%) (Fig. 2-II, Suppl. Fig. 2).

#### Morphotype III

Found in *Leptodactylus latrans, Leptodactylus validus, Boana platanera, Leptodactylus insularum, Leptodactylus fuscus, Leptodactylus fragilis, Scinax rostratus, Scinax ruber, Trachycephalus typhonius, Boana lanciformis, Rhinella sternosignata, Adenomera hylaedactyla, Rhinella horribilis*. Oval-shaped parasite (average length and width 29.02 µm ± 7.78 x 13.87 µm ± 4.09) with granular cytoplasm that usually presents 1-2 large vacuoles. It has a characteristic elongated nucleus (8.89 µm ± 2.59 x 2.34 µm ± 0.69) with two chromatin condensations at each end. The kinetoplast (6.34 µm ± 3.12) is posterior to the nucleus. It serves as the point of origin for a wavy membrane that extends outward and covers 40-60% of the parasite’s total length. The axoneme is visible, and the maximum width of the membrane is 1.37 µm ± 0.66. A free flagellum is usually observed and has an approximate length of 10.91µm ± 6.3 (Fig. 2-III, Suppl. Fig. 3).

#### Morphotype IV

Found in *Hypsiboas maculateralis, Boana pellucens, Scinax elaeochroa,* and *Smilisca phaeota*. Globular, with a bulging structure (L: 37.5 µm ± 14.1 ). Dark and utterly granular cytoplasm. The widest part of the parasite is delimited by a thin undulating membrane, with a marked axoneme and 6-8 distinct undulations. In most cases, it has a free flagellum roughly 4.10 µm ± 5.89 µm long. The nucleus is elongated (DN: 7.23 µm ± 2.98, dN: 3.16 µm ± 1.51), wholly condensed, and the kinetoplast is poorly observable (Fig. 2-IV, Suppl. Fig. 4).

#### Morphotype V

Found in *Lithodytes lineatus, Hypsiboas maculateralis*. It has an elongated U-shaped morphology (L: 66.59 µm ± 5.86, W: 10.34 µm ± 0.17). It has pointed edges at both the anterior and posterior ends. Its cytoplasm has evident longitudinal striations throughout the cell body. The nucleus is round or oval (maximum width 4.71 µm ± 0.30 x 4.24 µm ± 0.470) and is located in the central part of the parasite, leaning towards its exterior, adjacent to the undulating membrane. The kinetoplast (0.49 µm ± 0.13) is conspicuous and located towards the parasite’s posterior half. The undulating membrane emerges from the kinetoplast and goes to the anterior edge. Undulations are occasionally observed, and their number might vary between specimens (±7). Sometimes, it is possible to observe a very thin free flagellum with a length of approximately 16.58 µm ± 2.97 (Fig. 2-V, Suppl. Fig. 5).

#### Morphotype VI

Found in *Leptodactylus validus, Leptodactylus insularum,* and *Scinax aff. ruber. Trypanosoma* thin and elongated (L: 30.50 µm ± 0.35, W: 5.68 µm± 1.18 ), with a coiled arrangement (on itself or in an S shape). The cytoplasm is dark, lacking granulations or striations. The nucleus is oval (5.32 µm ± 0.38 x 2.74 µm ± 0.04), barely perceptible, occupying almost the width of the parasite. It is characterized by a thick, undulating membrane, approximately 1.88 µm ± 0.12, without a distinct axoneme. The membrane folds appear to envelop the entire perimeter of the parasite. The kinetoplast is circular, displaced towards the posterior end, and barely visible, along with its adjacent flagellar pocket (Fig. 2-VI, Suppl. Fig. 6).

#### Morphotype VII

Found in *Boana platanera, Leptodactylus latrans, Scinax aff. ruber, Leptodactylus fuscus,* and *Rhinella horribilis.* Parasite with an oval shape (L: 33.14 µm ± 8.01, W: 18.13 µm ± 6.43) and well-defined edges. Its cytoplasm is condensed, with fine granulations. The nucleus is elongated (8.97 µm ± 1.57 x 3.04 µm ± 0.16) and typically does not present condensations, although in some cases a condensation could be observed towards the anterior end. The kinetoplast is evident and is located adjacent to the posterior end of the nucleus. This morphotype is characterized by an undulating membrane extending from the kinetoplast throughout the parasite’s cytoplasm, forming 4-5 apparent C-shaped undulations. The free flagellum cannot be observed (Fig. 2-VII, Suppl. Fig. 7).

#### Morphotype VIII

Found in *Scinax aff. ruber*. Elongated vermiform parasite (L: 44.01 µm ± 1.13, W: 7.07 µm ± 0.67), U-shaped or slightly curved, with rounded edges and condensed cytoplasm. When visible, the nucleus is elongated (9.99 µm ± 1.35 × 2.29 µm ± 0.23 µm), and it has chromatin condensations at each end. Its kinetoplast is visible and is next to a distinguishable flagellar pocket. The morphotype is characterized by an undulating membrane that surrounds the entire body of the parasite, giving its cytoplasm a segmented appearance. No free flagellum was observed (Fig. 2-VIII, Suppl. Fig. 8).

#### Morphotype IX

Found in *Boana platanera, Hypsiboas maculateralis*. Small bullet-shaped *Trypanosoma* (L: 30.94 µm ± 15.01 x 5.99 µm ± 2.42), with a rounded posterior end and pointed anterior end. It has a free flagellum of approximately 9.46 µm ± 7.13. The cytoplasm is completely granular, presenting small refringent vacuoles. The nucleus is conspicuous and slightly oval (6.23 µm ± 3.27 x 2.15 µm ± 0.51). The undulating membrane extends throughout approximately 90% of the parasite’s total length. It has a visible axoneme and free flagellum with approximately 9.46 µm ± 7.13 in length (Fig. 2-IX, Suppl. Fig. 9).

#### Morphotype X

Found in *Lithobates palmipes*. Elongated and thin morphotype (L: 34.51 µm ± 2.70, W: 1.52 µm ± 0.09). It can be arranged in an S shape or with a slight curve. It has a somewhat oval nucleus (L: 2.58 µm ± 0.31, W: 1.30 µm ± 0.12) that is surrounded by two white vacuoles close to each of its ends. The undulating membrane is thin (0.79 µm ± 0.19), and the axoneme cannot be seen, nor its free flagellum (Fig. 2-X, Suppl. Fig. 10).

#### Morphotype XI

Found in *Scinax ruber*. It is an elongated parasite with rounded edges; its arrangement is flattened or or similar to an entangled network. (L: 40.77 µm ± 2.71, W: 14.18 µm ± 0.31). Non-observable kinetoplast and nucleus. Its main feature is its dense, granular cytoplasm, which often presents striations or wavy edges (Fig. 2-XI, Suppl. Fig. 11).

#### Morphotype XII

Found in *Lithobates vaillanti*. It is characterized by its elongated shape (L: 56.82 µm ± 1.42, W: 18.98 µm ± 0.44) and its pointed edges. It has longitudinal striations throughout the cytoplasm, along with numerous vacuoles. Its nucleus is round and conspicuous (4.38 µm ± 0,24 x 3.84 µm ± 0.11), and a more hyaline border can be observed around it. Its kinetoplast is smaller than the previous morphotypes (0.613 µm ± 0.013). The presence of approximately eight undulations on its membrane is a common feature of the specimens of this morphotype (Fig. 2-XII, Suppl. Fig. 12).

Among the morphotypes identified, it is worth highlighting that the first three were the most recurrent, with morphotype III present in 57.9% of the samples, morphotype II in 54.5% and morphotype I in 15.9%, while morphotypes X, XI, and XII were the least recurrent, present in 1-2% of the samples. It is essential to note that numerous co-infections exist among all morphotypes (Suppl. Table S3). Mixed infections were found in 48.9% of the samples (43/88).

**Fig 2.**
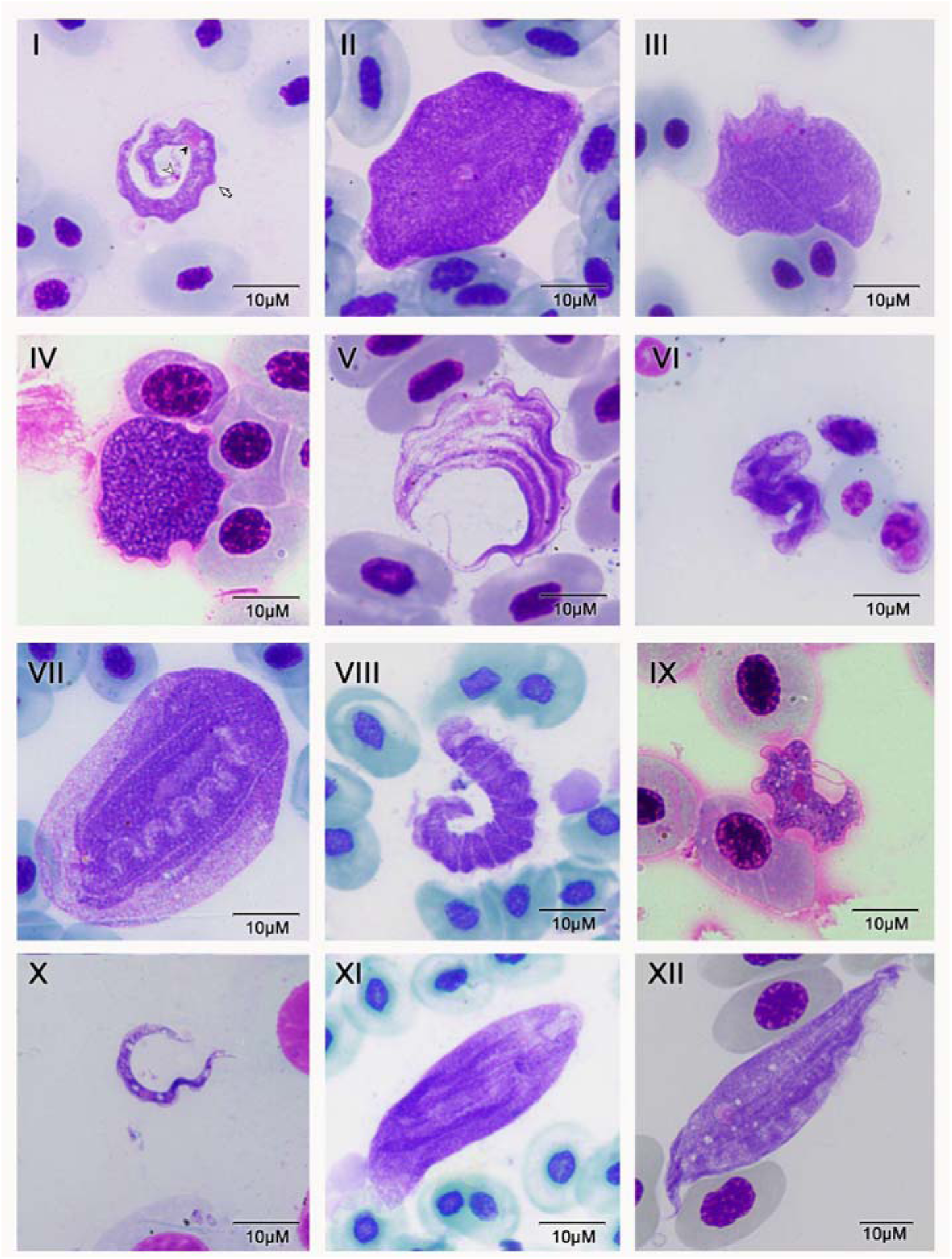
Illustrative photomicrographs of the twelve morphotypes identified in blood smears (Giemsa-stained) in Colombian anurans. Black arrows, short white arrows, and long white arrows indicate nuclei, kinetoplasts, and undulating membrane, respectively—scale bar: 10 μm.

### 3.3 Statistical analysis

All 12 morphotypes were measured, and their morphological and morphometrical features are shown in Table 1. Only three out of the twelve morphotypes showed a frequency of occurrence that allowed us to take more than 30 different photographs for the subsequent statistical analysis. Almost all variables did not have a normal distribution (p-value < 0.05 for all variables, except PN: posterior end to nucleus). However, Bispo and Marques (2023) report that a PCA is robust even under this condition.

The PCA showed two principal components: PC1 (body size vs. nuclear arrangement), which explained 64.35% of data variability, with higher loading values in the variables kinetoplast-nucleus (-0.343), width at nucleus level (0.332), posterior end to nucleus (-0.318), anterior end to nucleus (-0.316), and area (-0.316). In that way, larger parasites with a more central nucleus are located toward the right side of the graphic analysis (Morphotype II). In comparison, thinner parasites with a more displaced nucleus are located toward the left side of the graph (Morphotype I). On the other hand, PC2 (Nuclear shape vs undulating membrane width) explained 35.65% of data variability, with higher loading values in variables: primary nucleus diameter (-0.432), minor diameter of the nucleus (0.399), and undulating membrane width (-0.391) (Fig. 3a, Suppl. Table S4). Thus, parasites with a more rounded nucleus and thin membrane are located at the top (Morphotypes I and II), while parasites with elongated nuclei and a thick, undulating membrane are located at the bottom (Morphotype III) (Fig. 3a).

The ANOSIM analysis showed an all-group statistic of R=0.527 and p-value=0.0001; all comparisons between each pair of morphotypes showed p-values <0.0003 (Suppl. Table S5), indicating that the similarity within each morphotype is significantly higher than the similarity between different morphotypes. PERMANOVA analysis showed an all-group statistic of F=82.66 and p-value=0.0001; all comparisons between each pair of morphotypes showed p-values <0.0003 (Suppl. Table S5). The separation between groups was confirmed by the cladogram based on Gower distance (qualitative plus quantitative variables), where a clade for each of the three morphotypes was generated; the similarity between morphotypes II and III was higher than with the slimmer morphotype I (Fig. 3b).

**Fig 3.**
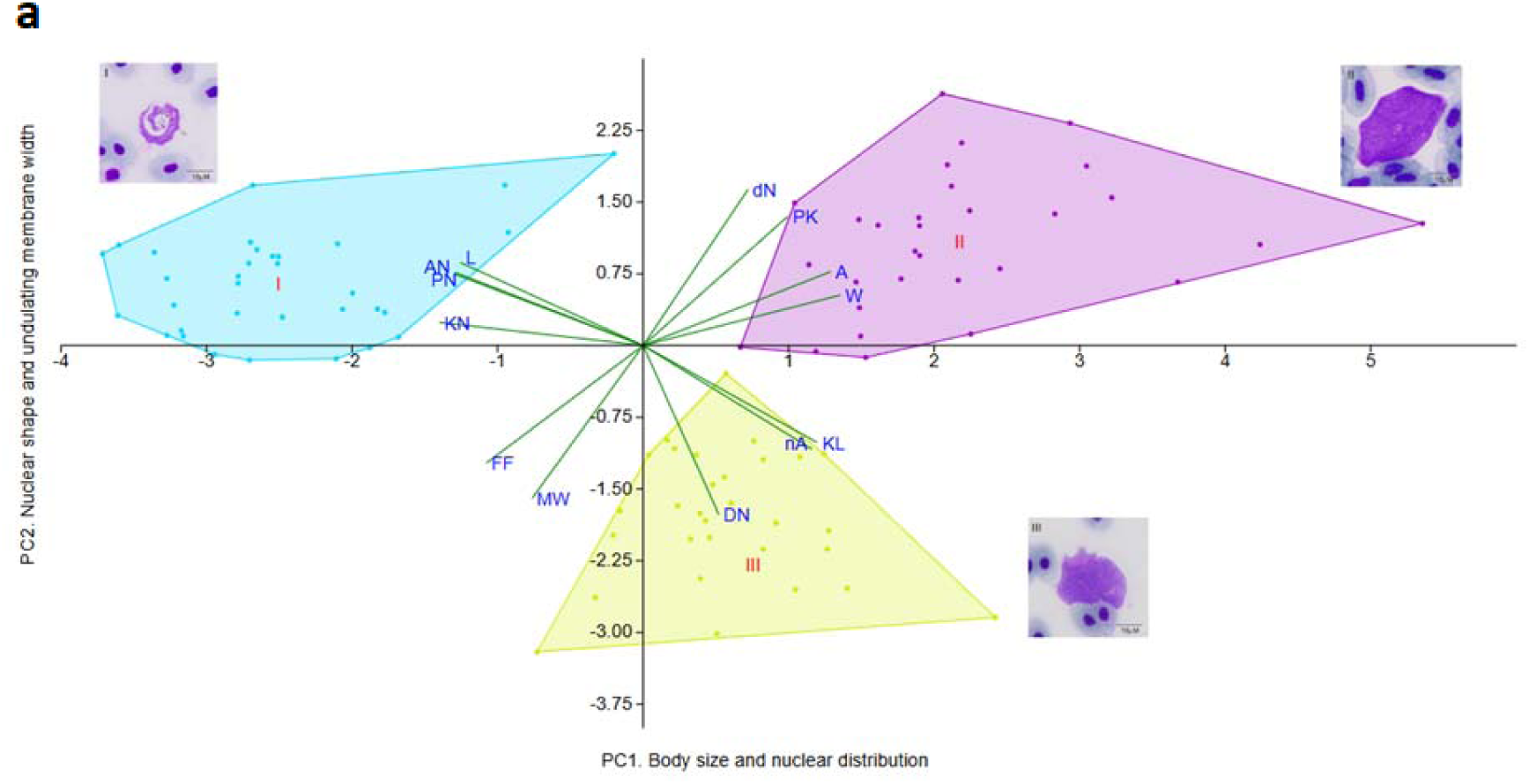
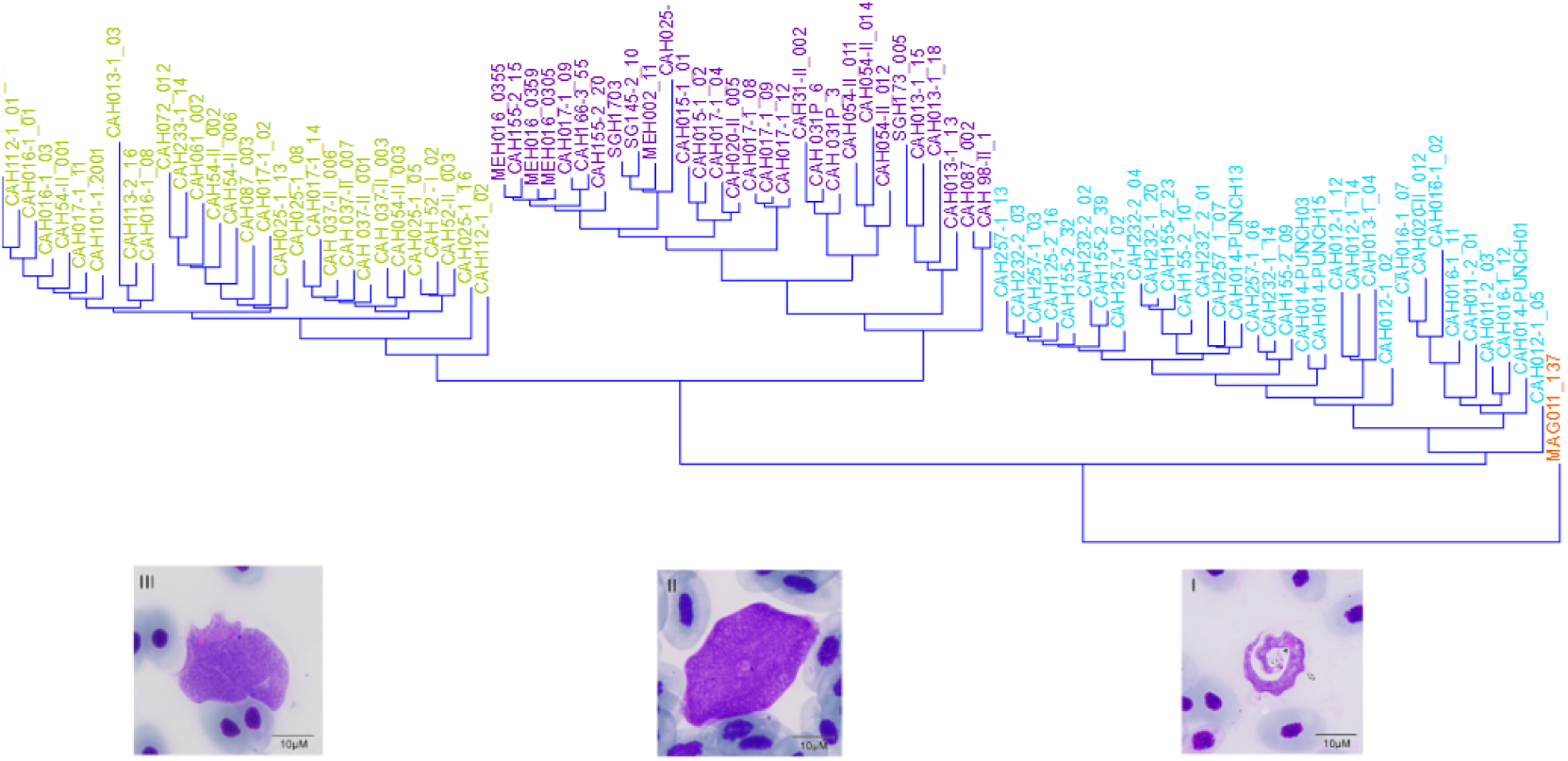
Morphotypes I (blue), II (purple), and III (yellow) grouping using qualitative and quantitative variables. **A.** Principal Component Analysis from morphometric measurements of morphotypes I, II, and III (n≥ 30 for each morphotype). **B.** Cladogram based on Gower distance of qualitative and quantitative variables described in Table 1. Morphotype XII (MAG011_137) was used as the root of the cladogram.

### 3.3 Phylogenetic analysis

Eight 18S sequences were obtained and deposited in GenBank. They originate from samples identified as single infections through microscopic analysis, which allowed us to associate them with four of the previously described morphotypes, as follows: Morphotype I, PQ860802 of *Leptodactylus latrans*. Morphotype II: PQ860796 of *Rhinella horribilis* and PV157257 from *Rhinella horribilis* culture. Morphotype III: PQ860800 and PQ860801 of *Boana platanera*, and PQ860798 of *Trachycephalus typhonius*. Morphotype IV: PV157255 from *Boana pellucens*. Morphotype V: PX285885 from *Lithodytes lineatus*.

The maximum likelihood phylogenetic tree is presented in Figure 4. This phylogeny has two clearly separated clades, with bootstrap values greater than 70; a clade containing *Trypanosoma* lineages from aquatic vertebrates and reptiles, and adjacent to it, a clade that comprises anurans’ *Trypanosomas.* This last one also includes several lineages isolated from vectors, as well as a reptile lineage (*Smaug depressus*).

**Figure 4.**
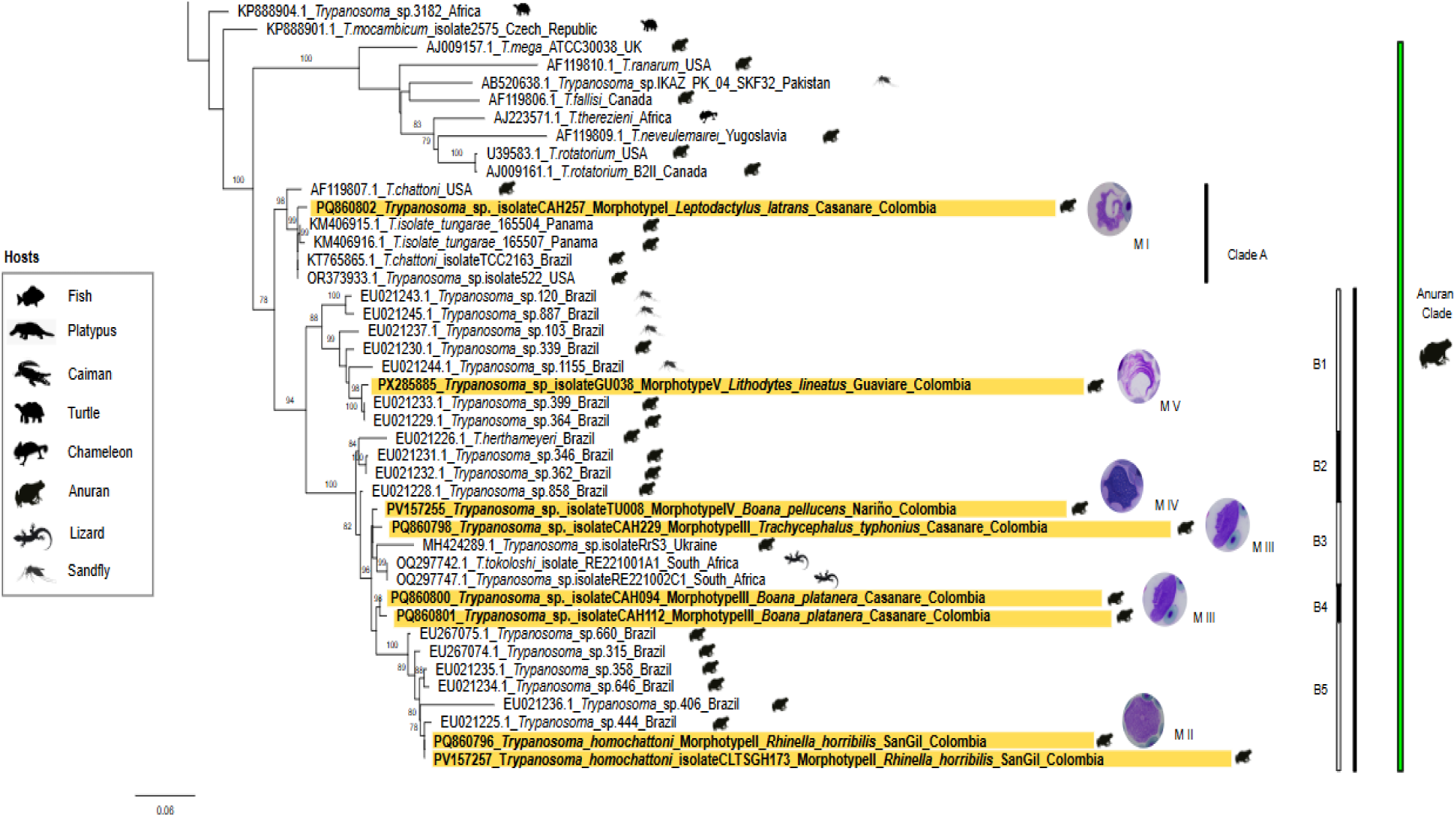
Zoom of maximum likelihood phylogenetic hypotheses based on 700 bp fragments for 18S RNA. This figure illustrates the relationships between different morphospecies and molecular lineages of *Trypanosoma* identified in fish, crocodiles, platypus, chameleons, and anurans (See Suppl. Fig. 14). Molecular lineages characteristic of morphotypes I, II, III, IV, and V are highlighted horizontally in yellow. The bootstrap value is indicated above each branch (See complete image Suppl Fig. 14).

The anuran clade is composed of two main clades (Figure 4). Clade A groups the morphospecies I together with *T. chattoni* and *T. tungarae* (Bernal & Pinto, 2016). Clade B is subdivided into five clades (B1 to B5). In Clades B1 and B2, parasites infecting anurans (i.e., *Sciopemyia* sp.) and their possible vectors (i.e., *Evandromyia infraspinosa*) have been identified (Ferreira et al., 2008; da S. Ferreira et al., 2015). Clade B1 includes lineages from Colombia and Brazil, revealing broader geographic and host associations. Within this clade, the lineage corresponding to morphotype V (PX285885), identified in *Lithodytes lineatus,* is also present. This lineage is closely related to lineage 399, previously reported in *Rhinella margaritifer*, and lineage 364 from *Rhinella marina*, both of which are in the same clade and originate from Brazil (Ferreira et al. 2008; da S. Ferreira et al. 2015). These findings suggest a possible evolutionary relationship among trypanosomes infecting phylogenetically and ecologically distinct anuran hosts across a wide geographical range.

Clades B3 and B4 appear to comprise a species complex, consisting of several closely related clades. Within these clades, morphospecies III and IV appear to be associated with Trypanosoma tokoloshi (OQ297742.1), a parasite found in a reptile (Smaug depressus) in South Africa, with genetic distances of 1.43–1.87% and 1.14%, respectively (see Supplementary Table 3). Ultimately, clade B5 is a well-supported clade in which Morphospecies II is grouped with Trypanosoma spp. from Brazil.

Some authors have found that a difference of more than 2% in 18S rRNA is used to separate wildlife trypanosome species (Guhl & Ramírez, 2011; Lai et al., 2008; Carnes et al., 2015; Vallejo, 1998). This criterion is met in the case of fish trypanosomes: The distance is 4.38% between *T. pleuronectidium* (DQ016613.1) and *T. boissoni* (U39580.1); 12% between *T. abeli* (KR052821.1) and T. murmanensis (DQ016616.1); and 8.41% between T. triglae (U39584.1) and *T. granulosum*_Portugal (AJ620552.1) (see Figure 4). The same is true for amphibian trypanosomes: for instance, the distance between *T. herthameyeri* (EU021226.1) and *T. rotatorium* (U39583.1) is 9.52%, while the distance between T. herthameyeri (EU021226.1) and *T. tokoloshi* (OQ297742.1) and 5.36% between *T. rotatorium_*USA (U39583.1) and *T. fallisi* (AF119806.1) (see Supplementary Table 3). Therefore, the genetic distance separating morphospecies II from its closest neighbours in the phylogeny enables us to propose it as a new species of *Trypanosoma* in amphibians.

Description of *Trypanosoma homochattoni* n. sp. (Morphotype II) (Suppl. Fig. 2).

This parasite is round or slightly oval in shape. Its cytoplasm is completely granular and may exhibit some striations toward the outer edges. Its length ranges from 32.86 µm ± 6.71 x 28.609 µm ± 5.52, and it covers a substantial area (730.61 µm ± 263.75). It is characterized by having a round, centrally located nucleus (PN: 15.41 µm ± 4.31, AN: 15.97 µm ± 3.00) and a kinetoplast that can be found above the nucleus, located toward one of its edges (KN: 4.79 ± 6.23). From the kinetoplast, a petite flagellum can be observed emerging diffusely.

Etymology: The species name reflects the morphological similarity of the parasite with *Trypanosoma chattoni* (Mathis and Leger, 1911).

Remarks:

Although *Trypanosoma homochattoni* n. sp. resembles *T. chattoni* (ATCC 50294 strain, AF119807.1) morphologically, the genetic distance between the two organisms is 7.22%, which supports its classification as a different species (Guhl & Ramírez, 2011; Lai et al., 2008; Carnes et al., 2015; Vallejo, 1998). *T. homochattoni* also tends to be larger than *T. chattoni,* with an amoeboid disposition and a more developed flagellum. *T. homochattoni,* found infecting amphibians in Colombia, is morphologically similar to morphotype M10 of Trypanosoma sp. 444 (EU021225.1; Ferreira et al., 2007), a parasite that infects *Leptodactylus chaquensis* in Brazil. The genetic distance between these two parasites is just 0.43%, suggesting that they may be the same species. Furthermore, *T. homochattoni* shares morphological similarities with *T. chattoni*, which infects *Leptodactylus fuscus* (Schneider, 1799) in Brazil (Lemos et al., 2008; Ferreira et al., 2008). The genetic distance with the parasite reported in *Rhinella horribilis* is 7%, which could indicate that this parasite should be renamed *T. homochattoni* n. sp. and that this species might have a broader distribution in South America. Regarding its distribution in Colombia, analysis of samples at the GERPH Collection identified *T. homochattoni* as a generalist parasite, as it was found in 14 anuran species from eight families across Colombia.

In the type sample UNAL: GERPH: SGH173, morphological variation in the length of the nucleus was observed, potentially indicating division processes occurring in the bloodstream (Jones and Woo, 1986; Souza, 2001). This new species has a generalist behavior, identified in 14 anuran species from eight distinct families across Colombia.

Taxonomy:

Phylum: Euglenozoa Cavalier-Smith, 1981

Class: Kinetoplastea Honigberg, 1963, emend. Vickerman,1976

Subclass: Metakinetoplastina Vickerman, 2004

Order: Trypanosomatida Kent, 1880

Family: Trypanosomatidae Doflein, 1951

Genus: *Trypanosoma* Gruby, 1843

Subgenus: *Trypanosoma* Gruby, 1843 emend. Votýpka and Kostygov, 2022

Type host: *Rhinella horribilis* (Wiegmann, 1833) (Bufonidae, *Rhinella*)

Representative DNA sequences: submitted to GenBank as 18S partial sequence PQ860796, PV157257.

Type locality: Colombia, Santander, San Gil (6.425° N, 73.201° W)

Type specimens: Hapantotype peripheral blood smear deposited in the Grupo Estudio Relación Parásito

Hospedero (GERPH) Collection UNAL: GERPH: SGH173 Parasitemia: 26 Trypanosomatids.

Site of infection: Peripheral blood

Vector: Unknown.

Additional hosts and additional material available on request from Biological Collection GERPH, Biology Department-Universidad Nacional de Colombia: *Leptodactylus latrans* (CAH013, CAH015, CAH017, CAH073, CAH087, CAH098, CAH118, CAH125, CAH155, CAH195, CAH232, CAH257)*, Leptodactylus validus* (CAH020)*, Scinax rostratus* (CAH031, CAH085, CAH101)*, Leptodactylus insularum* (CAH008, CAH044, CAH124)*, Boana platanera* (CAH025, CAH054, CAH113, MEH002, TIH002)*, Boana boans* (MEH003)*, Scinax aff. ruber* (CAH231)*, Trachycephalus typhonius* (CAH121, CAH122, CAH251)*, Leptodactylus colombiensis* (CAH160, MEH016)*, Lithobates palmipes* (MEH011)*, Smilisca phaeota* (TU010)*, Leptodactylus fuscus* (CAH065, CAH259)*, Hypsiboas maculateralis* (GU009).

Geographical distribution: Brazil, Pantanal, wetland central region, Mato Grosso do Sul, Miranda (Supplementary Table 4), Colombia: a) Santander, San Gil, b) Casanare, Yopal, Vereda Quebrada Seca, Hato Santa Teresa, c) Casanare, Trinidad, Vereda el Pozo petrolero, Finca Matepalma, d) Cundinamarca, Medina, Mesa de Cura, e) Cundinamarca, Tibacuy, Vereda El Mango, Finca Angel Ramos, f) Nariño, Tumaco, g) Guaviare, Sam Jose de Guaviare, Puentes naturales.

Ocurrence: 39/76. It was calculated as the number of samples with morphotype II (including single and coinfections) over the total number of positive samples determined by microscopy in this study.

## 4. Discussion

Neotropical countries harbour a great diversity of amphibians, hosting an under-explored but surely immense diversity of *Trypanosoma* species. In this study, we identified 12 distinct *Trypanosoma* morphotypes across 25 amphibian species, and we described *Trypanosoma homochattoni*, a new and generalist species found in 14 anuran species across Colombia.

We employed statistical methods that integrated both morphometric and categorical variables to distinguish these morphotypes. This approach enhances the accuracy of *Trypanosoma* identification in anuran blood samples, particularly in groups where morphological features are commonly used to differentiate between species but comprehensive taxonomic keys are lacking. Principal Component Analysis (PCA) revealed clear separation between morphotypes I, II, and III based on body size, nuclear arrangement, and undulating membrane width, accounting for 64.35% of the total variance. Nuclear and membrane measurements are commonly employed in morphometric analyses (Jones & Woo, 1986; Woo, 1969; Lemos et al., 2008; Spodareva et al., 2018). However, this study suggests that the total area of the parasite could also be a critical quantitative variable to consider.

Additionally, Gower’s similitude measurement, appropriate for mixed-source data, further supported the differentiation between morphotypes and enhanced the characterization of these parasites. The integration of morphometric and categorical data was essential for distinguishing morphotypes in co-infected samples.

We detected that Morphotype I shows a strong resemblance to *T. tungarae,* reported by Bernal and Pinto (2016). Similarly, Morphotype II shares features with *T. chattoni*, and Morphotype III is similar to *Trypanosoma* sp. (Tipo 3), previously identified in *Craugastor fitzingeri* and *Atelopus longirostris* in Colombia (Carvajal, 1982). While these three morphotypes display considerable overlapping in size and other features, some differences, such as the shape of the nucleus and undulating membrane, are evident.

By utilizing integrative taxonomy, we identified five morphotypes with their respective barcode sequences: Morphotype I (lineage PQ860802), Morphotype II (lineage PQ860796), Morphotype III (lineages PQ860800, PQ860801, and PQ860798), Morphotype IV (lineage PV157255), and Morphotype V (lineage PX285885). These findings supported the geographical expansion of *T. tungarae*, previously reported in Panama (Bernal & Pinto, 2016), and confirmed its presence in *Leptodactylus latrans* from Casanare, Colombia. Furthermore, we identified the *T. homochattoni n.* sp., which morphologically resembled *T. chattoni,* but they show a genetic distance in the molecular marker 18S rRNA.

We also identified Morphotype V, which resembles *T. fallisi* and *Trypanosoma* sp. (Type 4) reported by Carvajal (1982) in Colombia. Morphotype VII shares characteristics with *T. herthameyeri* and *T. rotatorium*. A noteworthy observation was the identification of Morphotype VIII, which was found only in *Scinax ruber* and has not been previously reported. Its distinctive and conserved features suggest it may represent a novel *Trypanosoma* species. However, confirmation of this hypothesis requires further research, including barcode sequencing from a single infection.

Morphotype IX displays a fusiform body and a granularity pattern similar to a morphotype reported in *Amietia delalandii* from Africa (Du Buisson, 2023). Morphotype X, which was rarely observed, exhibits an unusual morphology more closely resembling fish trypanosomes such as *Trypanosoma pseudobagri* (Gu et al., 2007), *T. carassii* (Overath et al., 1998), and other morphotypes from fish and leeches in Brazil (Lemos et al., 2015). These findings suggest potential cross-taxon transmission, possibly through shared aquatic vectors (Hayes et al., 2014; Attias et al., 2016; Spodareva et al., 2018).

Morphotype XII, detected at low frequency in *Lithobates vaillanti* from Magdalena (Suppl. Table S3), shows distinctive features such as longitudinal striations, a circular nucleus with a hyaline border, and numerous undulations that resemble *Trypanosoma* sp. (Tipo 9) from *Lithobates palmipes* in Valle del Cauca (Carvajal, 1982) and the R1 morphotype from *Lithobates clamitans* in the USA (Shannon and Bolek, 2023). Despite originating from different regions, all hosts belong to the Ranidae family, suggesting a possible family-level specificity.

While morphotypes II and III were frequently found, morphotypes X and XII appear more host-specific. In contrast, morphotypes IV, VI, and XI infect multiple species but do not resemble any previously reported forms, indicating these may represent first-time records.

The genetic analysis also revealed patterns that reinforce the idea of species complexes, particularly between Morphotypes III and IV, which exhibited variable morphologies but shared similar molecular lineages. For instance, Morphotype III (lineages PQ860800, PQ860801, and PQ860798) and Morphotype IV (lineage PV157255) exhibited genetic distances of approximately 1.4%, 1.9%, 1.4%, and 1.1%, respectively, from *Trypanosoma tokoloshi* (OQ297742.1). This phenomenon mirrors the *T. theileri* complex, where closely related lineages infect different hosts of cattle and cervids (Filip-Hutsch et al., 2022). The relatively high genetic distances observed among these morphotypes further highlight the complexity of *Trypanosoma* taxonomy, especially when morphological similarities can obscure underlying genetic differentiation.

Moreover, *T. tokoloshi*, which primarily infects reptiles, showed some molecular similarities with Morphotype IV. It had a genetic distance of 9–10%, suggesting potential overlap between amphibian and reptilian *Trypanosoma* species. These genetic similarities and distinct morphologies indicate the need for further phylogenetic studies to clarify these relationships. The genetic distances found among *Trypanosoma* morphotypes in this study support the existence of multiple, genetically distinct lineages not easily differentiated by morphology alone.

One key challenge in the field remains the high pleomorphism of these parasites, combined with frequent co-infections, which make species-level identification difficult. Although molecular techniques offer a potential solution, they are often hindered by the presence of nucleated blood cells, which complicate DNA extraction and sequencing (Ferreira et al., 2008; Megía-Palma et al., 2024). Furthermore, the reliance on 18S rRNA data, both in this study and globally, presents a limitation for comparative genetic analyses. Incorporating additional molecular markers could strengthen the molecular framework and help confirm or refute the hypotheses generated from 18S-based analyses. (Hamilton et al., 2004; Nishimoto et al., 2008).

*Leptodactylidae* frogs exhibited a particularly high prevalence of infection, consistent with previous observations in Brazil, where species from the *Hylidae, Leptodactylidae,* and *Bufonidae* were most commonly infected (Leal et al., 2009). The behavior of *Leptodactylidae* species, such as digging deep burrows, might facilitate parasite dispersal, particularly in regions with high amphibian density during the dry season (Oliveira & Giaretta, 2008; Ponssa & Medina, 2016). Further studies should focus on unraveling the host-parasite relationships, especially considering the geographic distribution of these parasites and their potential transmission vectors, which may include leeches that are common to both aquatic and terrestrial environments.

It is worth highlighting that anurans occupy an intermediate evolutionary position between aquatic and terrestrial vertebrates (Bardsley & Harmsen, 2017). As a result, several hypotheses have been proposed, suggesting that amphibian trypanosomatids are historically older and that, through various evolutionary processes, the effect of host switching was amplified (Bartlett et al., 2009). This allowed these parasites to adapt to new terrestrial hosts. One factor that may drive the evolutionary processes between taxonomic groups is the presence of a common vector, such as leeches, which can infect both aquatic and terrestrial vertebrates (Spodareva et al., 2018). This evolutionary flexibility may explain why some morphotypes are more frequent or adapt more easily to specific hosts.

The impact of *Trypanosoma* infections on amphibian populations remains poorly understood. While some species, such as *Atelopus longirostris*, are critically endangered and infected with *Trypanosoma*, the consequences of these infections on their health and conservation status remain unknown (Carvajal, 1982). As such, future research should explore the ecological and evolutionary dynamics of these parasitic infections, particularly their impact on host fitness and potential roles in amphibian declines.

## 5. Conclusions

This study demonstrates substantial trypanosome diversity among Colombian amphibians, characterized by pronounced morphological variation. The application of statistical analyses, including principal component analysis (PCA) and permutational multivariate analysis of variance (PERMANOVA), in conjunction with standardized molecular techniques, facilitated the identification of genetic lineages corresponding to five distinct morphotypes. Coinfections were observed in 43.4% of cases. Phylogenetic analyses corroborated established host-parasite associations and identified novel relationships, such as an expanded distribution for *T. tungarae* and the identification of a new species, *T. homochattoni.* Multiple lineages were found to be related to the *T. tokoloshi* species complex. To further refine trypanosome taxonomy and elucidate infection dynamics, future research should incorporate host ecological data, vector identification, and advanced molecular methodologies.

## Acknowledgments

We thank Andres David Jimenez Maldonado, Martha Calderon, Gustavo Andrés Fuentes, and Ana Saldarriaga for collecting and sampling some of the specimens used in this study. We thank Juanita Cárdenas Sánchez, Alejandra Quiroz, Maria Fernanda Medina, Brayan Gamboa, and Karly Ojeda for their assistance with the handling and examination of blood slides.

## Ethics Declaration

The biological samples used in this study were obtained from specimens previously deposited in the GERPH biological collection. Only specimens of *Rhinella horribilis* were captured and manipulated for the purpose of obtaining positive controls. This procedure was authorized by Act No. 09-2023 of the Facultad de Ciencias, Universidad Nacional de Colombia. No other species were captured, handled, or explicitly sacrificed for this research. All procedures adhered to the relevant institutional and national guidelines for the ethical use of biological material.

## Consent to Publish declaration and Consent to Participate declaration

**not applicable.**

## Data Availability Statement

All data supporting the findings of this study are available within the article and its Supplementary Information files, including the complete database of samples, variable descriptions with graphical examples, additional statistical test data, and photographic plates for each morphotype.

## Authors contributions

N.E.M. conceived, designed the study, and provided the necessary resources. All five authors (A.T.O.R., A.D.G.G., A.F.A.G., C.M.V.L., and N.E.M.) participated in data curation and formal analysis, with A.T.O.R. playing a leading role in the latter. Funding acquisition was carried out by A.D.G.G., A.F.A.G., and N.E.M. Validation was performed by A.T.O.R., A.D.G.G., A.F.A.G., and C.M.V.L. All authors contributed to the investigation. Visualization was primarily conducted by C.M.V.L., with support from A.F.A.G. Methodology was developed by A.T.O.R., A.D.G.G., A.F.A.G., and N.E.M. N.E.M. supervised and administered the project. All authors were involved in the writing, review, and final approval of the manuscript.

## Declaration of generative AI and AI-assisted technologies in the writing process

During the preparation of this work, the author(s) (Angelica Ospina and Nubia E. Matta) used ChatGPT and Grammarly Service to improve the accuracy of the language and readability. After using this tool/service, the author(s) (Angelica Ospina and Nubia E. Matta) reviewed and edited the content as needed and take full responsibility for the content of the publication. All the authors reviewed and approved the final version.

