## Supplementary material for "Untangling the diversity of Trypanosomes infecting Colombian amphibians: morphometric and molecular insights"

Suppl. Table S1. Database with geographic and taxonomic information of the total number of samples analyzed. [.xlsx file]

Suppl. Table S2. Qualitative features with their respective assigned categories for the morphological description.

| Qualitative feature | Categories | Image with a related example |
|---|---|---|
| Shape | 1- Elongated | Suppl. figure 1A |
| | 2- Oval | Suppl. figure 2D |
| | 3- Ameboid | Suppl. figure 2B |
| Arrangement | 1- Rolled | Suppl. figure 1C |
| | 2- Crushed | Suppl. figure 2C |
| | 3- Extended | Suppl. figure 2A |
| | 4- S-shaped | Suppl. figure 6D |
| | 5- Concave or slightly curved | Suppl. figure 9A |
| | 6- Undulating | Suppl. figure 10B |
| Anterior border shape | 1- Pointed | Suppl. figure 10D |
| | 2- Rounded | Suppl. figure 7B |
| | 3- Blunt | Suppl. figure 3B |
| | 0- n/a | Suppl. figure 2C |
| Posterior border shape | 1- Pointed | Suppl. figure 1B |
| | 2- Rounded | Suppl. figure 9D |
| | 3- Blunt | Suppl. figure 6D |
| | 0- n/a | Suppl. figure 2D |
| Cytoplasm texture | 1- Striated | Suppl. figure 5B |
| | 2- Granular | Suppl. figure 4D |
| | 3- Mixed | Suppl. figure 7C |
| | 4- Granular up to the core | Suppl. figure 1A |
| | 5- Smooth | Suppl. figure 6C |
| Vacuoles | 1- Yes | Suppl. figure 4A |
| | 2- No | Suppl. figure 5A |
| | 3- Yes, around the core | Suppl. figure 1C |
| Core shape | 1- Rounded | Suppl. figure 5D |
| | 2- Oval | Suppl. figure 3A |
| | 3- Elongated | Suppl. figure 7A |
| Kinetoplast shape | 1- Circular | Suppl. figure 12D |
| | 2- Oval | Suppl. figure 5B |
| Undulating membrane | 1- Yes, undulating | Suppl. figure 4A |

| | | |
|---|---|---|
| | 2- Yes, flattened | Suppl. figure 7C |
| | 3- No | Suppl. figure 2D |
| Visible axoneme | 1- Yes | Suppl. figure 4B |
| | 2- No | Suppl. figure 8C |
| Number of ridges | Set number (e.g. 6) | Suppl. figure 5D |

Suppl. Table S3. Amphibian species positive for trypanosoma by microscopy, according to the twelve morphotypes found, together with parasitemia and location data.

| **Sample ID (GERPH collection)** | **Accession number** | **Locality in Colombia** | **Family** | **Host species** | **Morphotype determined** | **Parasitaemia** |
|---|---|---|---|---|---|---|
| GERPH:CAH008 | | Casanare | Leptodactylidae | *Leptodactylus insularum* | II, III | 1 |
| GERPH:CAH009 | | Casanare | Leptodactylidae | *Leptodactylus latrans* | I | 1 |
| GERPH:CAH011 | | Casanare | Leptodactylidae | *Leptodactylus latrans* | I | 2 |
| GERPH:CAH012 | | Casanare | Leptodactylidae | *Leptodactylus latrans* | I,III | 6 |
| GERPH:CAH013 | | Casanare | Leptodactylidae | *Leptodactylus latrans* | I, II, III | 13 |
| GERPH:CAH014 | | Casanare | Leptodactylidae | *Leptodactylus latrans* | I, III | 15 |
| GERPH:CAH015 | | Casanare | Leptodactylidae | *Leptodactylus latrans* | II | 2 |
| GERPH:CAH016 | | Casanare | Leptodactylidae | *Leptodactylus latrans* | I, III | 9 |
| GERPH:CAH017 | | Casanare | Leptodactylidae | *Leptodactylus latrans* | II, III | 1 |
| GERPH:CAH018 | | Casanare | Leptodactylidae | *Leptodactylus latrans* | III | 4 |
| GERPH:CAH019 | | Casanare | Leptodactylidae | *Leptodactylus validus* | III, VI | 6 |
| GERPH:CAH020 | | Casanare | Leptodactylidae | *Leptodactylus validus* | I, II | 5 |
| GERPH:CAH021 | | Casanare | Leptodactylidae | *Leptodactylus validus* | II, III | 1 |
| GERPH:CAH025 | | Casanare | Hylidae | *Boana platanera* | II,III,VII | 4 |
| GERPH:CAH028 | | Casanare | Leptodactylidae | *Leptodactylus validus* | - | 1 |
| GERPH:CAH029 | | Casanare | Hylidae | *Scinax rostratus* | - | 1 |
| GERPH:CAH031 | | Casanare | Hylidae | *Scinax rostratus* | II, III | 1 |
| GERPH:CAH037 | | Casanare | Hylidae | *Boana platanera* | III | 5 |
| GERPH:CAH043 | | Casanare | Phyllomedusidae | *Pithecopus hypochondrialis* | II | 1 |
| GERPH:CAH044 | | Casanare | Leptodactylidae | *Leptodactylus insularum* | II, III | 6 |

| | | | | | | |
|---|---|---|---|---|---|---|
| GERPH:CAH052 | | Casanare | Hylidae | *Boana platanera* | III, IX | 3 |
| GERPH:CAH053 | | Casanare | Hylidae | *Boana platanera* | III, IX | 1 |
| GERPH:CAH054 | | Casanare | Hylidae | *Boana platanera* | II, III, VII, IX | 5 |
| GERPH:CAH058 | | Casanare | Hylidae | *Boana platanera* | VII | 1 |
| GERPH:CAH059 | | Casanare | Hylidae | *Boana platanera* | III | 4 |
| GERPH:CAH061 | | Casanare | Leptodactylidae | *Leptodactylus fuscus* | III | 1 |
| GERPH:CAH065 | | Casanare | Leptodactylidae | *Leptodactylus fuscus* | II | 1 |
| GERPH:CAH067 | | Casanare | Leptodactylidae | *Leptodactylus fragilis* | III | 1 |
| GERPH:CAH072 | | Casanare | Leptodactylidae | *Leptodactylus fragilis* | III | 7 |
| GERPH:CAH073 | | Casanare | Leptodactylidae | *Leptodactylus latrans* | II | 3 |
| GERPH:CAH085 | | Casanare | Hylidae | *Scinax rostratus* | II | 1 |
| GERPH:CAH087 | | Casanare | Leptodactylidae | *Leptodactylus latrans* | II, III | 2 |
| GERPH:CAH094 | PQ860800 | Casanare | Hylidae | *Boana platanera* | III | 1 |
| GERPH:CAH097 | | Casanare | Leptodactylidae | *Leptodactylus insularum* | III | 1 |
| GERPH:CAH098 | | Casanare | Leptodactylidae | *Leptodactylus latrans* | I, II | 2 |
| GERPH:CAH100 | | Casanare | Leptodactylidae | *Leptodactylus insularum* | III | 1 |
| GERPH:CAH101 | | Casanare | Hylidae | *Scinax rostratus* | II, III | 3 |
| GERPH:CAH102 | | Casanare | Ranidae | *Lithodytes lineatus* | V | 3 |
| GERPH:CAH112 | PQ860801 | Casanare | Hylidae | *Boana platanera* | III | 5 |
| GERPH:CAH113 | | Casanare | Hylidae | *Boana platanera* | II, III | 19 |
| GERPH:CAH114 | | Casanare | Hylidae | *Scinax ruber* | II, III | 7 |
| GERPH:CAH118 | | Casanare | Leptodactylidae | *Leptodactylus latrans* | II | 1 |
| GERPH:CAH119 | | Casanare | Leptodactylidae | *Leptodactylus validus* | - | 1 |
| GERPH:CAH121 | | Casanare | Hylidae | *Trachycephalus typhonius* | II, III | 27 |
| GERPH:CAH122 | | Casanare | Hylidae | *Trachycephalus typhonius* | II, III, VII | 41 |
| GERPH:CAH124 | | Casanare | Leptodactylidae | *Leptodactylus insularum* | II | 2 |
| GERPH:CAH125 | | Casanare | Leptodactylidae | *Leptodactylus latrans* | I, II, III | 23 |
| GERPH:CAH126 | | Casanare | Hylidae | *Boana platanera* | III | 1 |
| GERPH:CAH141 | | Casanare | Leptodactylidae | *Leptodactylus latrans* | I | 3 |
| GERPH:CAH154 | | Casanare | Leptodactylidae | *Leptodactylus insularum* | VI | 1 |

| | | | | | | |
|---|---|---|---|---|---|---|
| GERPH:CAH155 | | Casanare | Leptodactylidae | *Leptodactylus latrans* | I, II, III, VII | 4 |
| GERPH:CAH160 | | Casanare | Leptodactylidae | *Leptodactylus colombiensis* | II | 1 |
| GERPH:CAH166 | | Casanare | Hylidae | *Scinax ruber* | II, III, VI, VII, VIII, XI | 118 |
| GERPH:CAH195 | | Casanare | Leptodactylidae | *Leptodactylus latrans* | II | 2 |
| GERPH:CAH209 | | Casanare | Hylidae | *Scinax ruber* | VII | 1 |
| GERPH:CAH212 | | Casanare | Hylidae | *Scinax rostratus* | III | 3 |
| GERPH:CAH216 | | Casanare | Leptodactylidae | *Leptodactylus insularum* | III | 3 |
| GERPH:CAH228 | | Casanare | Hylidae | *Scinax rostratus* | III | 4 |
| GERPH:CAH229 | PQ860798 | Casanare | Hylidae | *Trachycephalus typhonius* | III | 14 |
| GERPH:CAH231 | | Casanare | Hylidae | *Scinax aff. ruber* | II, VII | 184 |
| GERPH:CAH232 | | Casanare | Leptodactylidae | *Leptodactylus latrans* | I, II | 2 |
| GERPH:CAH233 | | Casanare | Hylidae | *Scinax ruber* | III, VII, VIII, XI | 32 |
| GERPH:CAH251 | | Casanare | Hylidae | *Trachycephalus typhonius* | II, III | 32 |
| GERPH:CAH256 | | Casanare | Leptodactylidae | *Leptodactylus* | II | 1 |
| GERPH:CAH257 | PQ860802 | Casanare | Leptodactylidae | *Leptodactylus latrans* | I | 15 |
| GERPH:CAH259 | | Casanare | Leptodactylidae | *Leptodactylus fuscus* | II, III, VII | 5 |
| GERPH:GU009 | | Guaviare | Hylidae | *Hypsiboas maculateralis* | II, IV, V, IX | 14 |
| GERPH:GU038 | | Guaviare | Ranidae | *Lithodytes lineatus* | V | 1 |
| GERPH:MAG005 | PX285885 | Magdalena | Ranidae | *Lithobates vaillanti* | II, XII | 3 |
| GERPH:MAG011 | | Magdalena | Ranidae | *Lithobates vaillanti* | II, XII | 18 |
| GERPH:MAG015 | | Magdalena | Ranidae | *Lithobates vaillanti* | II, XII | 1 |
| GERPH:MEH001 | | Cundinamarca - Medina | Hylidae | *Boana lanciformis* | III | 1 |
| GERPH:MEH002 | | Cundinamarca - Medina | Hylidae | *Boana platanera* | II, III | 35 |
| GERPH:MEH003 | | Cundinamarca - Medina | Hylidae | *Boana boans* | II | 84 |
| GERPH:MEH004 | | Cundinamarca - Medina | Bufonidae | *Rhinella sternosignata* | III | 6 |
| GERPH:MEH011 | | Cundinamarca - Medina | Ranidae | *Lithobates palmipes* | II, X | 16 |
| GERPH:MEH016 | | Cundinamarca - Medina | Leptodactylidae | *Leptodactylus colombiensis* | II, III | 21 |
| GERPH:MEH022 | | Cundinamarca - Medina | Craugastoridae | *Pristimantis sp.* | I | 1 |

| | | | | | | |
|---|---|---|---|---|---|---|
| GERPH:MEH059 | | Cundinamarca - Medina | Leptodactylidae | *Adenomera hylaedactyla* | III | 5 |
| GERPH:SGH145 | | Santander | Bufonidae | *Rhinella horribilis* | II, III, VII | 1 |
| GERPH:SGH148 | | Santander | Bufonidae | *Rhinella horribilis* | II, III, VII | 6 |
| GERPH:SGH172 | | Santander | Bufonidae | *Rhinella horribilis* | II, III | 2 |
| CLT_SGH173 | PQ860796, PV157257 (Culture) | Santander | Bufonidae | *Rhinella horribilis* | II | 26 |
| GERPH:TIH01 | | Cundinamarca - Tibacuy | Bufonidae | *Rhinella horribilis* | II, III | 2 |
| GERPH:TIH02 | | Cundinamarca - Tibacuy | Hylidae | *Boana platanera* | II, III | 3 |
| GERPH:TU008 | PV157255 | Nariño | Hylidae | *Boana pellucens* | IV | 8 |
| GERPH:TU009 | | Nariño | Hylidae | *Scinax elaeochroa* | IV | 7 |
| GERPH:TU010 | | Nariño | Hylidae | *Smilisca phaeota* | II, IV | 44 |

Suppl. Table S4. Loading values of the PCA statistical analysis performed on the data obtained from measurements of morphotypes I, II, and III

| PC | Eigenvalue | % variance |
|---|---|---|

| | | |
|---|---|---|
| 1 | 8.36593 | 64.353 |
| 2 | 4.63407 | 35.647 |
| **Loadings** | | |
| | PC 1 | PC 2 |
| L | -0.30748 | 0.2124 |
| W | 0.33208 | 0.12925 |
| A | 0.31575 | 0.18924 |
| AN | -0.31639 | 0.18729 |
| PN | -0.31775 | 0.18308 |
| DN | 0.12708 | -0.43202 |
| dN | 0.17651 | 0.39943 |
| nA | 0.28391 | -0.2651 |
| KL | 0.29239 | -0.2479 |
| KN | -0.34291 | 0.059235 |
| PK | 0.24322 | 0.33015 |
| MW | -0.18663 | -0.39104 |
| FF | -0.26351 | -0.30073 |

Suppl. Table S5. Representative values of the ANOSIM and PERMANOVA statistical analyses performed on the data obtained from measurements of morphotypes I, II, and III

| **ANOSIM** | | | **PERMANOVA** | |
|---|---|---|---|---|
| Permutation N: | 9999 | | Permutation N: | 9999 |

| R: | | 0.5268 | |
|---|---|---|---|
| p (same): | | 0.0001 | |
| **R-values upper diagonal<br>Bonferroni corrected the p-values lower diagonal** | | | |
| | I | II | III |
| I | | 0.7876 | 0.4249 |
| II | 0.0003 | | 0.4308 |
| III | 0.0003 | 0.0003 | |

| F: | | 82.66 | |
|---|---|---|---|
| p (same): | | 0.0001 | |
| **F values upper diagonal<br>Bonferroni corrected the p-values lower diagonal.** | | | |
| | I | II | III |
| I | | 141.5 | 33.35 |
| II | 0.0003 | | 50.7 |
| III | 0.0003 | 0.0003 | |

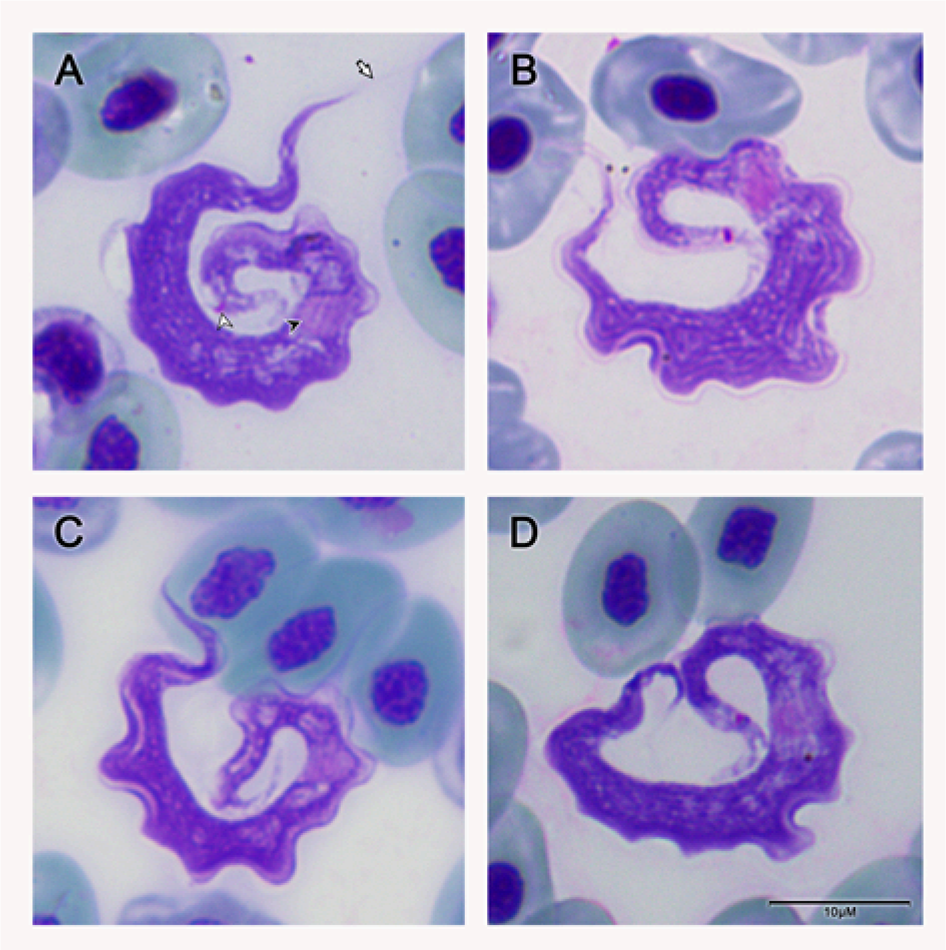

**Suppl. Fig. 1** Photomicrographs of morphotype I trypomastigotes (smears stained with 4% Giemsa). Black arrow, short white arrow, and long white arrow indicate nuclei, kinetoplast, and undulating membrane, respectively. Scale bar: 10 µm.

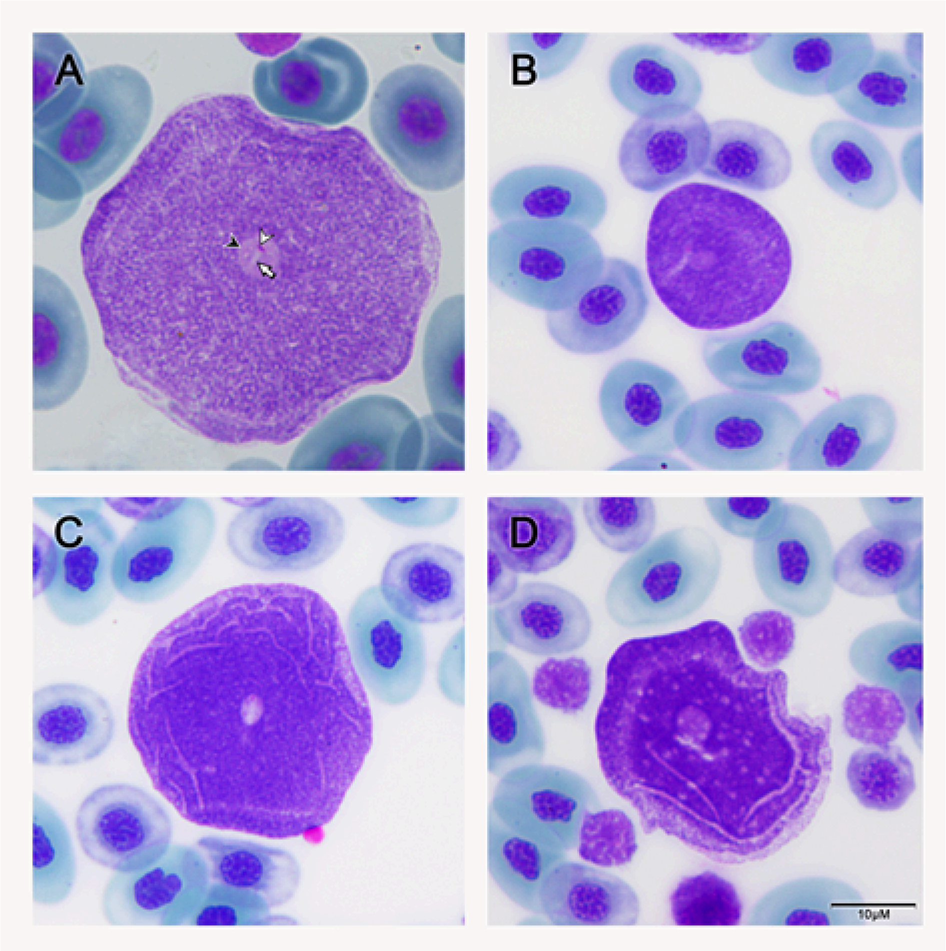

**Suppl. Fig. 2** Photomicrographs of morphotype II (smears stained with 4% Giemsa). Black arrow, short white arrow, and long white arrow indicate nuclei, kinetoplast, and undulating membrane, respectively. Scale bar: 10 μm.

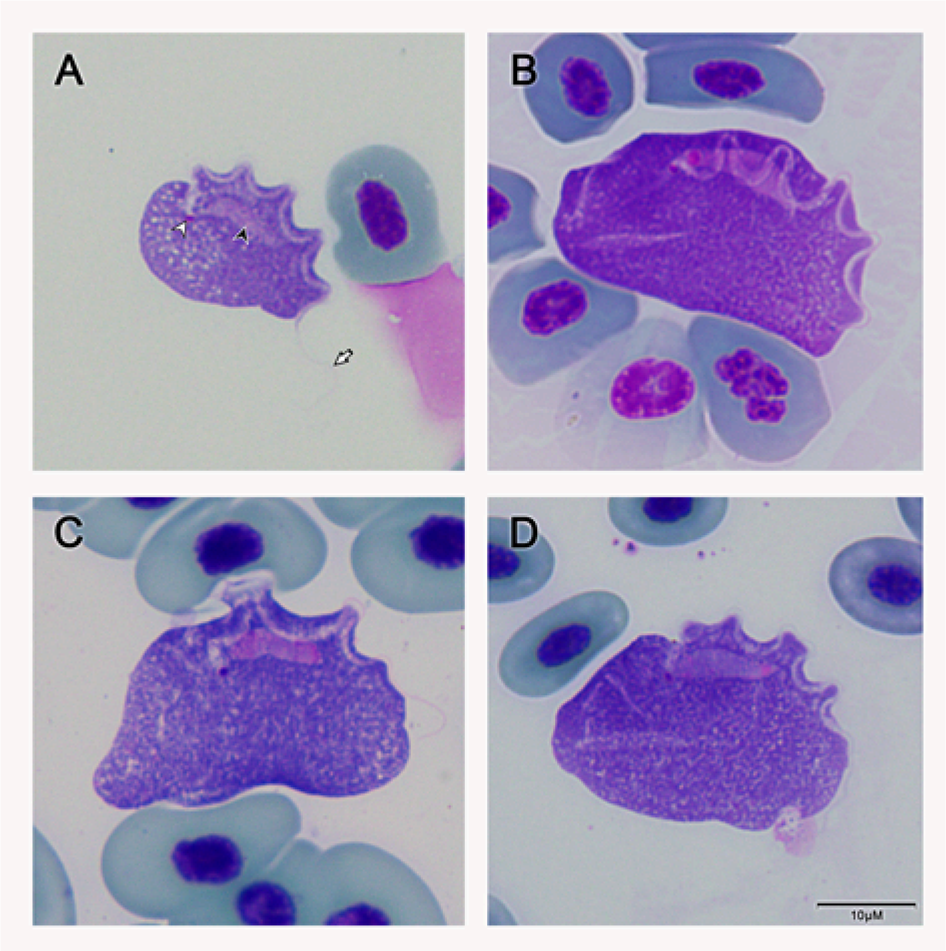

**Suppl. Fig. 3** Photomicrographs of morphotype III trypomastigotes (smears stained with 4% Giemsa). Black arrow, short white arrow, and long white arrow indicate nuclei, kinetoplast, and undulating membrane, respectively. Scale bar: 10 µm.

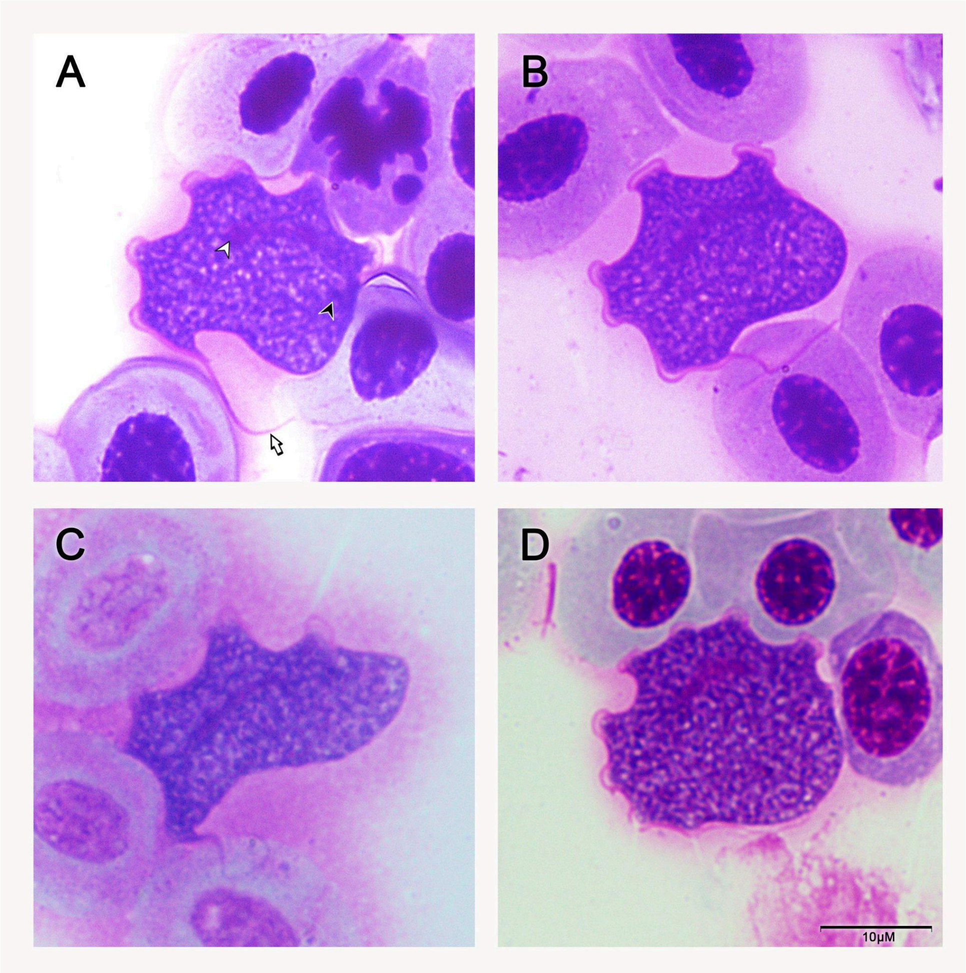

**Suppl. Fig. 4** Photomicrographs of morphotype IV trypomastigotes (smears stained with 4% Giemsa). Black arrow, short white arrow, and long white arrow indicate kinetoplast, nuclei and undulating membrane, respectively. Scale bar: 10 µm.

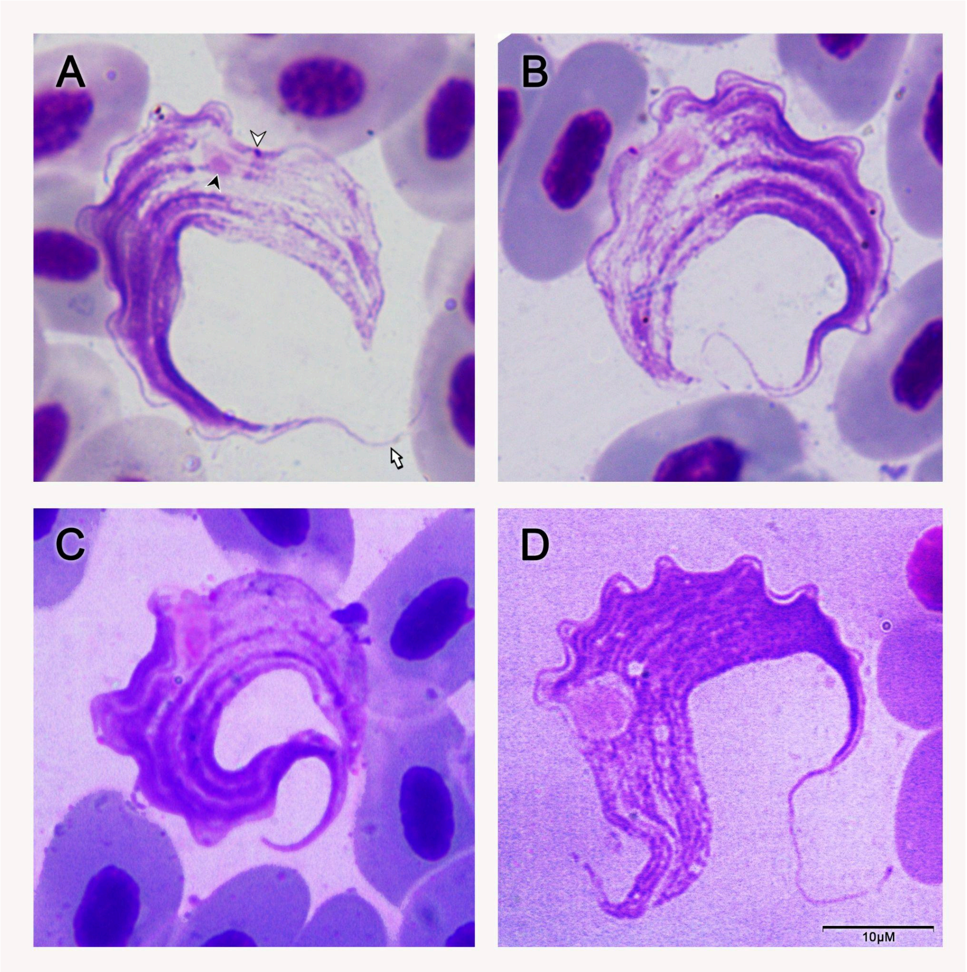

**Suppl. Fig. 5** Photomicrographs of morphotype V trypomastigotes (smears stained with 4% Giemsa). Black arrow, short white arrow, and long white arrow indicate nuclei, kinetoplast, and undulating membrane, respectively. Scale bar: 10 µm.

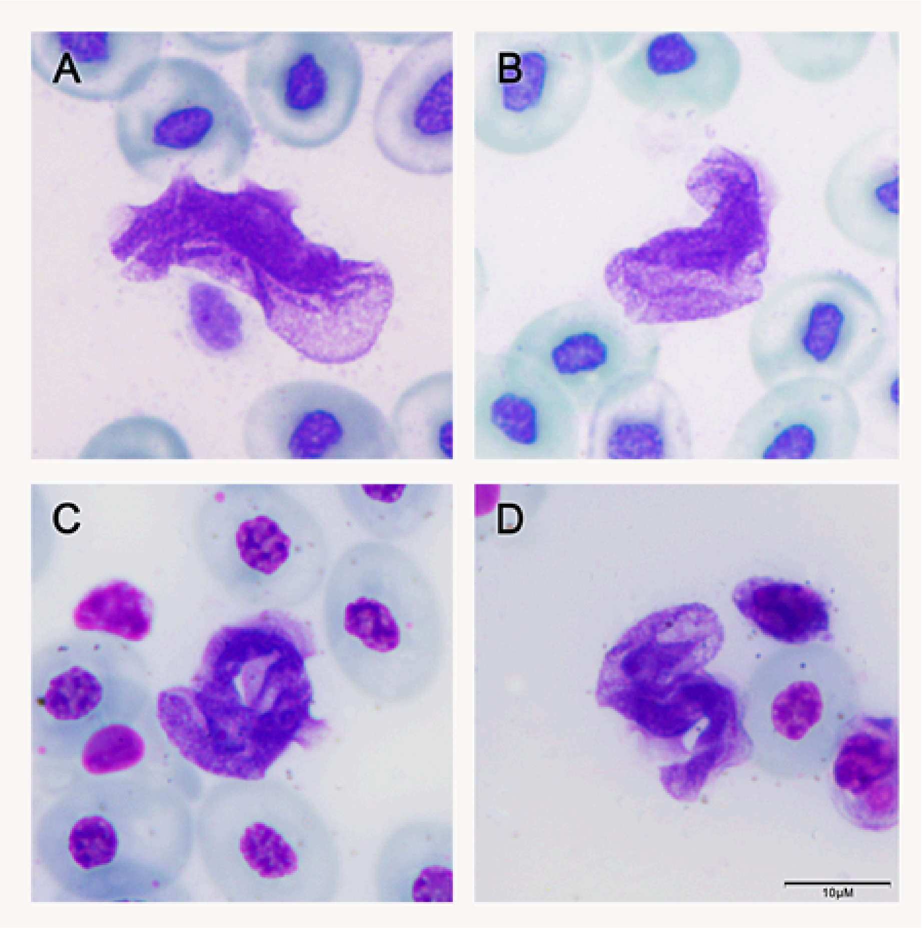

**Suppl. Fig. 6** Photomicrographs of morphotype VI trypomastigotes (smears stained with 4% Giemsa). Black arrow, short white arrow, and long white arrow indicate nuclei, kinetoplast, and undulating membrane, respectively. Scale bar: 10 µm.

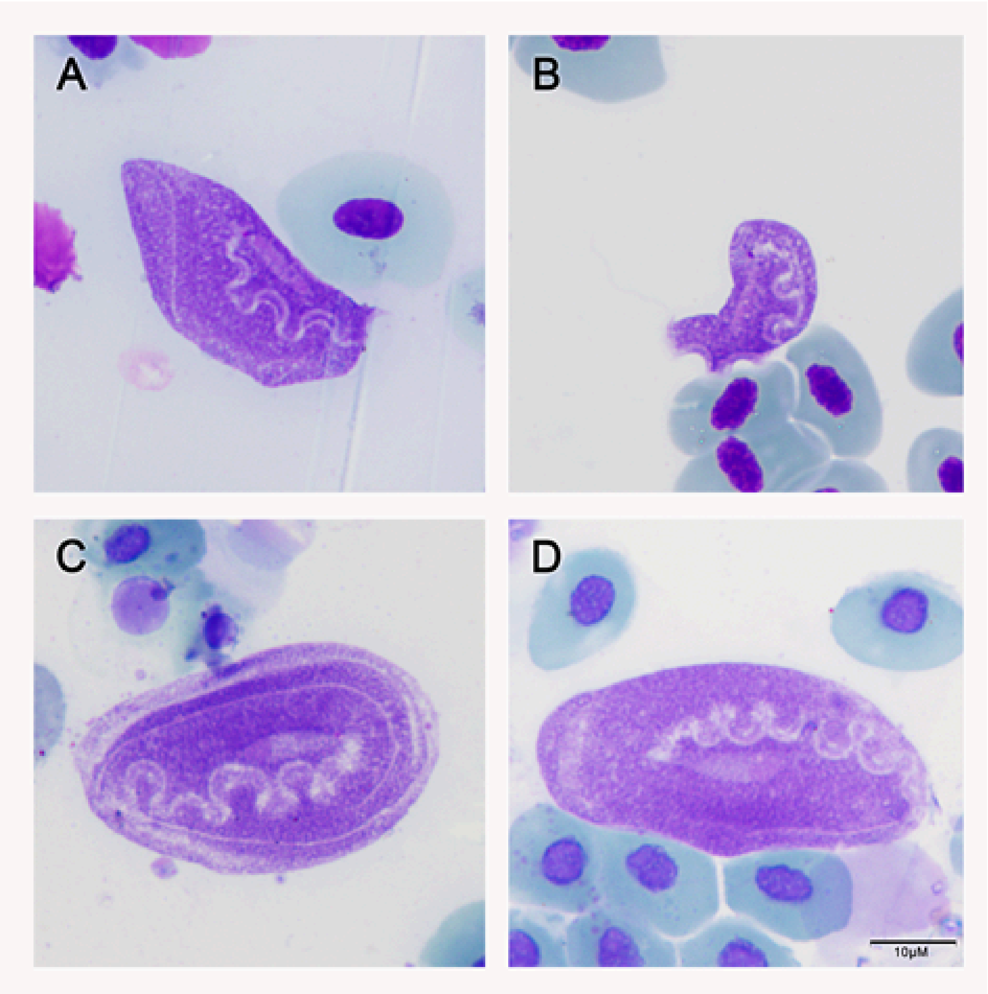

**Suppl. Fig. 7** Photomicrographs of morphotype VII trypomastigotes (smears stained with 4% Giemsa). Scale bar: 10 µm.

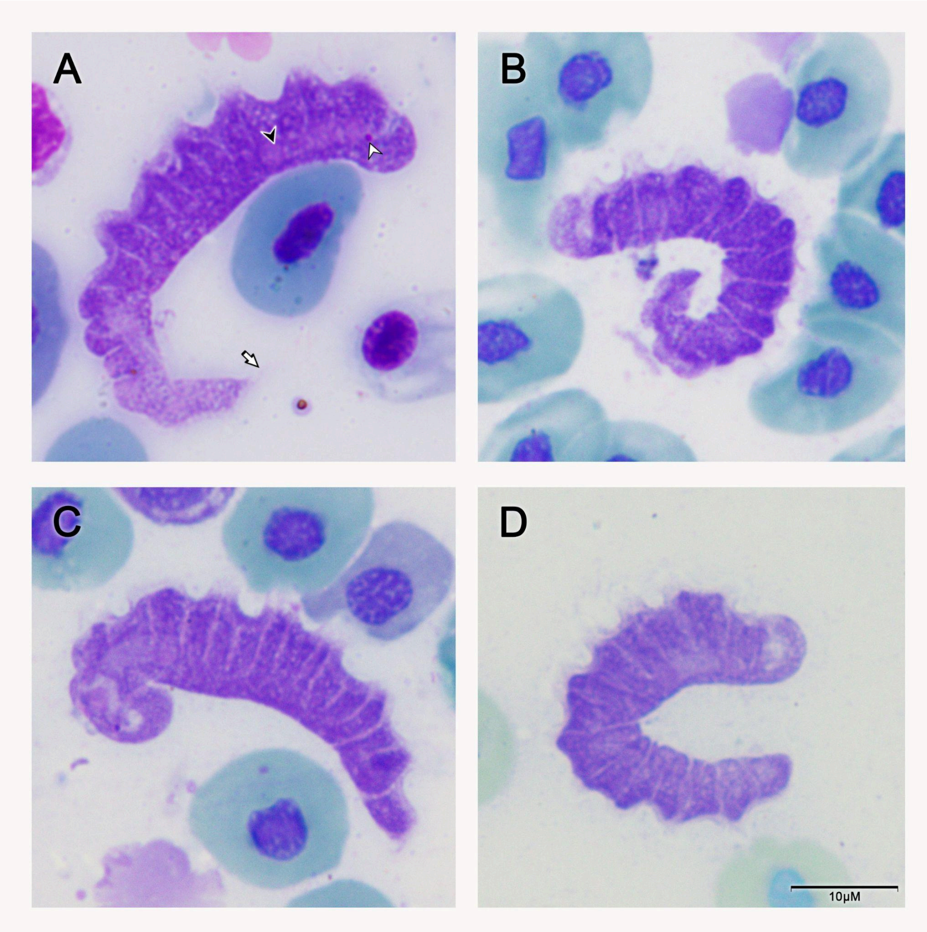

**Suppl. Fig. 8** Photomicrographs of morphotype VIII trypomastigotes (smears stained with 4% Giemsa). The short black arrow points to the nucleus, the short white arrow to the kinetoplast, and the long white arrow to the undulating membrane. The bar corresponds to 10 µm.

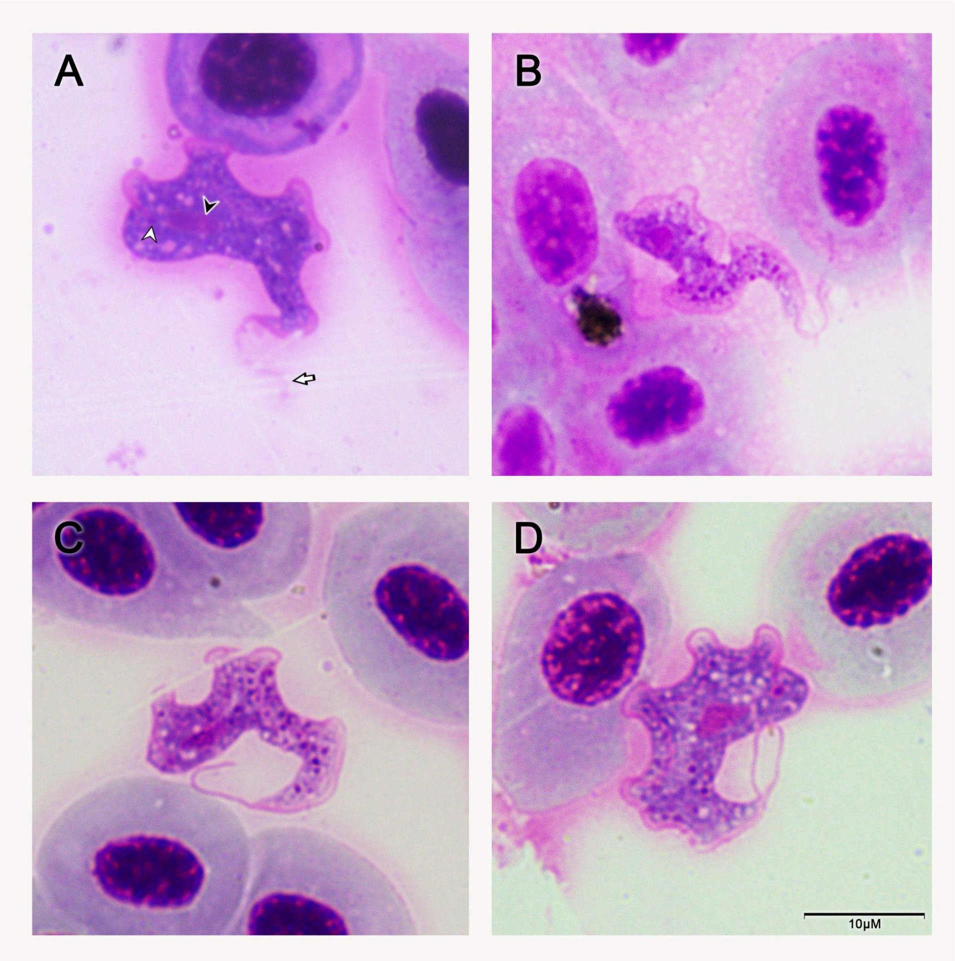

**Suppl. Fig. 9** Photomicrographs of morphotype IX trypomastigotes (smears stained with 4% Giemsa). The short black arrow points to the nucleus, the short white arrow to the kinetoplast, and the long white arrow to the undulating membrane. The bar corresponds to 10 µm.

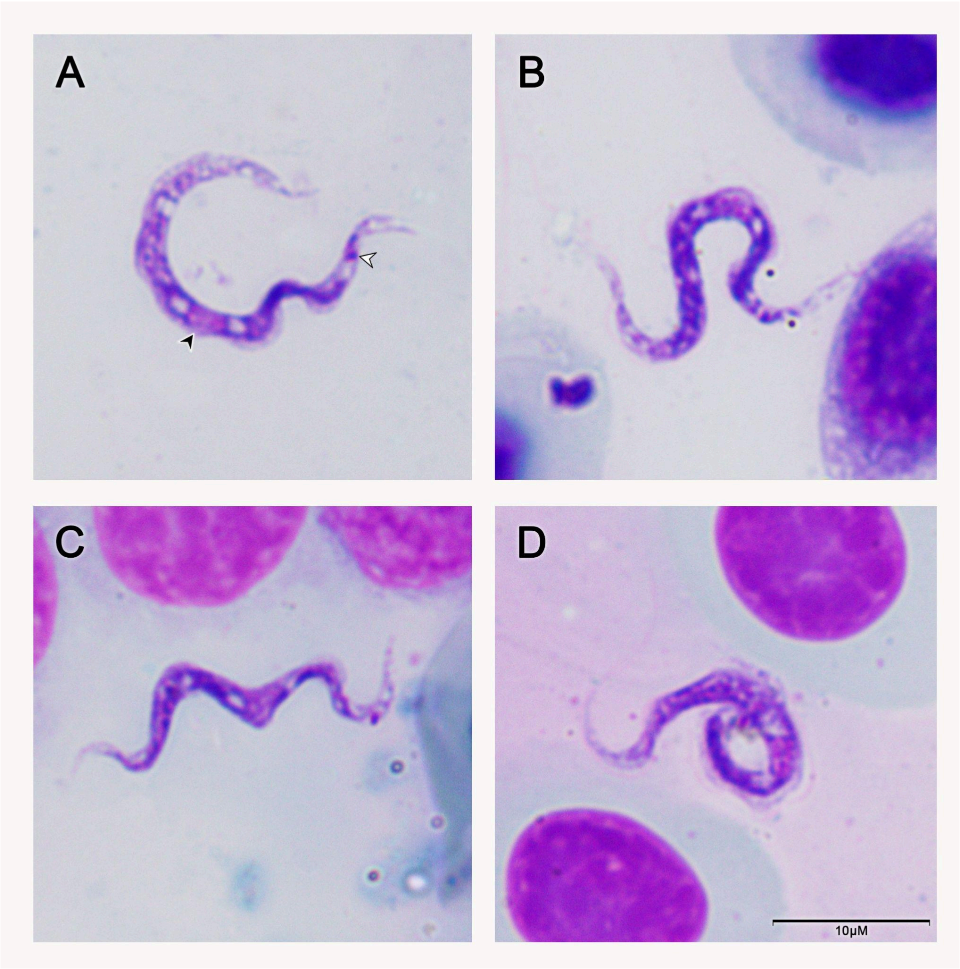

**Suppl. Fig. 10** Photomicrographs of morphotype X trypomastigotes (smears stained with 4% Giemsa). The short black arrow points to the nucleus and the short white arrow to the kinetoplast. The bar corresponds to 10 µm.

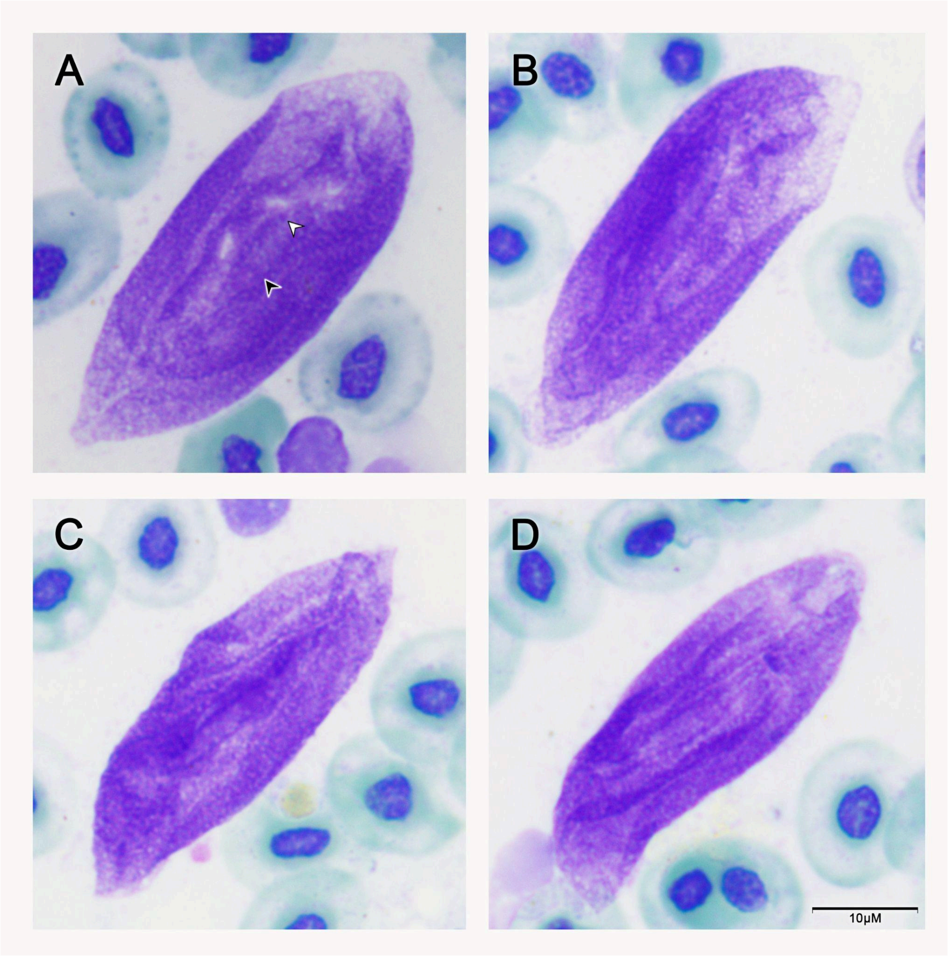

**Suppl. Fig. 11** Photomicrographs of morphotype XI trypomastigotes (smears stained with 4% Giemsa). The short black arrow points to the nucleus and the short white arrow to the kinetoplast. The bar corresponds to 10 µm.

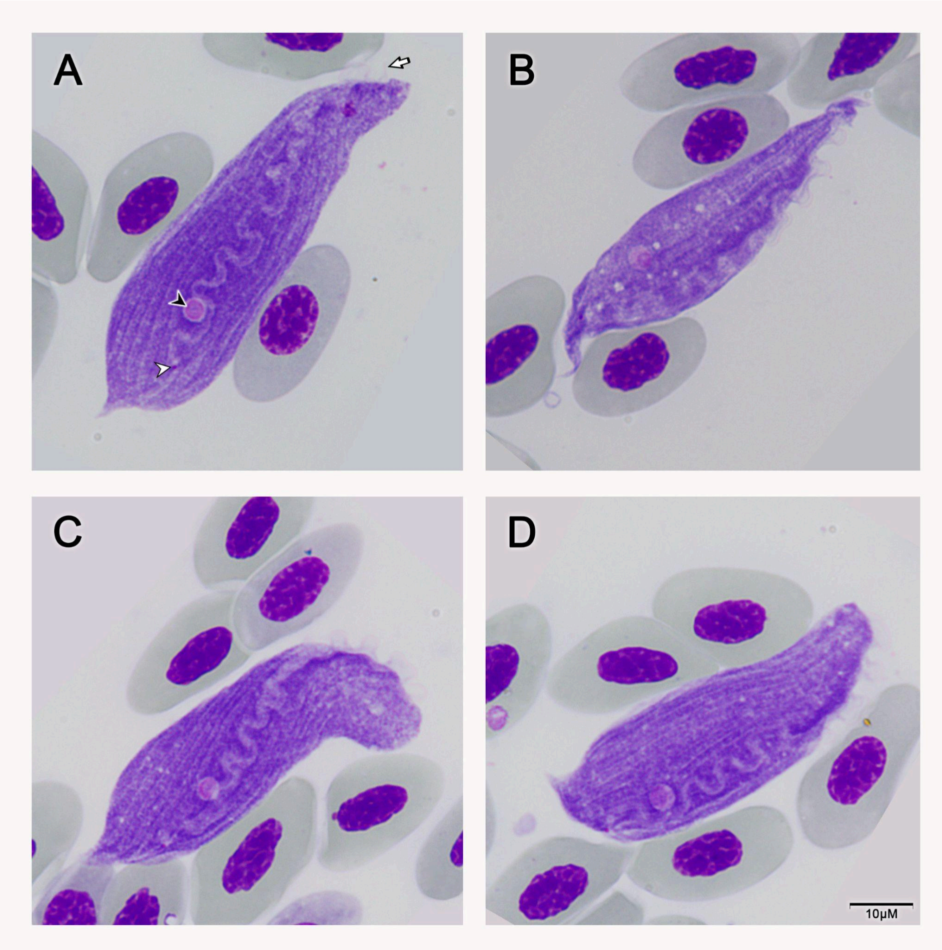

**Suppl. Fig. 12** Photomicrographs of morphotype XII trypomastigotes (smears stained with 4% Giemsa). The short black arrow points to the nucleus, the short white arrow to the kinetoplast, and the long white arrow to the undulating membrane. The bar corresponds to 10 µm.

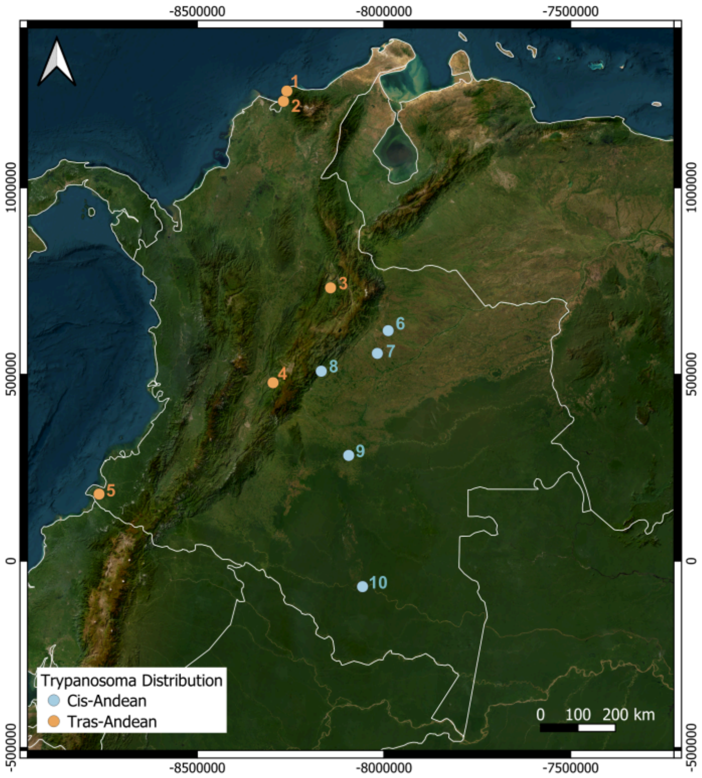

**Suppl. Fig. 13** Topographic map of Colombia created from satellite imagery and manually modified in QGIS (Geographic Information System) using WGS84 coordinates. The image indicates the sampled locations in Colombia (7 Departments, 10 sampling localities). The points indicate in the Trans-Andean area: 1. Magdalena (Santa Marta), 2. Magdalena (Pueblo Viejo), 3. Santander (San Gil), 4. Cundinamarca (Tibacuy), 5. Nariño (Tumaco). In the Cis-Andean area: 6. Casanare (Trinidad), 7. Casanare (El Yopal), 8. Cundinamarca (Medina), 9. Guaviare (San José del Guaviare), and 10. Caquetá (Solano).

**Suppl. Fig. 14 [SVG File].** Maximum likelihood phylogenetic hypotheses based on 700 bp fragments for 18S RNA. This figure illustrates the relationships between different morphospecies and molecular lineages of *Trypanosoma* identified in fish, crocodiles, platypus, chameleons, and anurans. Molecular lineages characteristic of morphotypes I, II, III, IV, and V are highlighted horizontally in yellow. The bootstrap value is indicated above each branch.
